# Systemic interindividual epigenetic variation in humans is associated with transposable elements and under strong genetic control

**DOI:** 10.1101/2022.05.27.493722

**Authors:** Chathura J. Gunasekara, Harry MacKay, C. Anthony Scott, Shaobo Li, Eleonora Laritsky, Maria S. Baker, Sandra L. Grimm, Goo Jun, Yumei Li, Rui Chen, Joseph L. Wiemels, Cristian Coarfa, Robert A. Waterland

## Abstract

Genetic variants can modulate phenotypic outcomes via epigenetic intermediates, for example by affecting DNA methylation at CpG dinucleotides (methylation quantitative trait loci – mQTL). Here, we present the first large-scale assessment of mQTL at human genomic regions selected for interindividual variation in CpG methylation (correlated regions of systemic interindividual variation – CoRSIVs). We used target-capture bisulfite sequencing to assess DNA methylation at 4,086 CoRSIVs in multiple tissues from 188 donors in the NIH Genotype-Tissue Expression (GTEx) program (807 samples total). At CoRSIVs, as expected, DNA methylation in peripheral blood correlates with methylation and gene expression in internal organs. We also discovered unprecedented mQTL at these regions. Genetic influences on CoRSIV methylation are extremely strong (median R^2^=0.76), cumulatively comprising over 70-fold more human mQTL than detected in the most powerful previous study. Moreover, mQTL beta coefficients at CoRSIVs are highly skewed (i.e., the major allele predicts higher methylation). Both surprising findings were independently validated in a cohort of 47 non-GTEx individuals. Genomic regions flanking CoRSIVs show long-range enrichments for LINE-1 and LTR transposable elements; the skewed beta coefficients may therefore reflect evolutionary selection of genetic variants that promote their methylation and silencing. Analyses of GWAS summary statistics show that mQTL polymorphisms at CoRSIVs are associated with metabolic and other classes of disease. A focus on systemic interindividual epigenetic variants, clearly enhanced in mQTL content, should likewise benefit studies attempting to link human epigenetic variation to risk of disease. Our CoRSIV-capture reagents are commercially available from Agilent Technologies, Inc.

**Significance Statement:** Population epigeneticists have relied almost exclusively on CpG methylation arrays manufactured by Illumina. At most of the >400,000 CpG sites covered by those arrays, however, methylation does not vary appreciably between individuals. We previously identified genomic loci that exhibit systemic (i.e. not tissue-specific) interindividual variation in DNA methylation (CoRSIVs). These can be assayed in blood DNA and, unlike tissue-specific epigenetic variants, do not reflect interindividual variation in cellular composition. Here, studying just 4,086 CoRSIVs in multiple tissues of 188 individuals, we detect much stronger genetic influences on DNA methylation (mQTL) than ever before reported. Because interindividual epigenetic variation is essential for not only mQTL detection, but also for epigenetic epidemiology, our results indicate a major opportunity to advance this field.

## Introduction

Genome-wide association studies (GWAS) have revolutionized the field of genetics by identifying genetic variants associated with a range of diseases and phenotypes (1–3). Nearly twenty years into the GWAS era, however, most human disease risk and phenotypic variation remain unexplained by common genetic variants (2), fueling interest in the possibility that individual epigenetic variation is an important determinant of phenotype (4, 5). To test this, over the last decade myriad studies have performed genome-scale screens to identify genomic regions at which epigenetic variation is associated with disease. Nearly all these epigenome-wide association studies (EWAS) used commercial arrays manufactured by Illumina (predominantly the HM450 and subsequently the scaled-up EPIC850 array) to assess methylation at CpG dinucleotides (a highly stable epigenetic mark) in peripheral blood DNA (6, 7). EWAS have uncovered associations between blood DNA methylation and neurological outcomes including Alzheimer’s disease (8), neurodegenerative disorders (9), educational attainment (10), and psychiatric diseases (11). The HM450 and EPIC arrays were instrumental in discoveries in epigenetic aging (12–14), smoking-induced DNA methylation alterations (15), and understanding how maternal smoking (16) and alcohol consumption (17) affect DNA methylation in newborns. Peripheral blood DNA methylation has been associated with birthweight (18), and body mass index (19).

The Illumina methylation arrays have also played a central role in advancing our understanding of genetic influences on CpG methylation. Genetic variants that correlate with methylation at a specific CpG site (usually in cis) are known as methylation quantitative trait loci (mQTL). Seminal observations of familial clustering of CpG methylation levels (20) led to the first formal study of mQTL (21), which utilized an early version of the Illumina methylation platform. Now, hundreds of studies, nearly all using Illumina methylation arrays, have investigated mQTL in humans (22), enabling estimates of methylation heritability and insights into how genetic effects on disease risk may be mediated by DNA methylation (23) and mechanisms of trans (inter-chromosomal) mQTL effects (24).

Despite these successes, existing and legacy Illumina methylation platforms are not ideal for population epigenetics. The success of GWAS was built upon the HapMap (25) and 1,000 Genomes (26) projects, which systematically mapped out human genome sequence variants so they could be assessed at the population level. So far, however, no ‘EpiHapMap’ project has been conducted. Several large consortium projects, including the Roadmap Epigenome Project (27), the Blueprint Epigenome Project (28), and the International Human Epigenome Consortium (29), focused primarily on characterizing tissue- and cell type-specific epigenetic variation rather than mapping out human genomic regions of interindividual epigenetic variation. The EWAS field therefore relied almost exclusively on Illumina arrays (30) which were designed without consideration of interindividual variation in DNA methylation (31, 32) and generally target CpGs that show little (33–36). To address this lacuna, we recently conducted an unbiased screen for correlated regions of systemic (i.e. not tissue-specific) interindividual epigenetic variation (CoRSIVs) in the human genome (37). Because that screen was based on only ten individuals, we set out to assess these regions in a larger cohort to characterize associations among interindividual genetic, epigenetic, and transcriptional variation. In addition to validating CoRSIVs as systemic epigenetic variants, assessing correlations with gene expression, and characterizing associations with transposable elements, we discovered that CoRSIVs exhibit much stronger mQTL than previously observed. Because interindividual variation is essential not just for mQTL detection but also for epigenetic epidemiology, our results have important implications for the EWAS field.

## Results

### Target-capture bisulfite sequencing confirms systemic interindividual variation in DNA methylation

In collaboration with the NIH Genotype-Tissue Expression (GTEx) program (38), we conducted target-capture bisulfite sequencing to quantify DNA methylation at 4,641 gene-associated CoRSIVs in multiple tissues representing the three embryonic germ layers from each of 188 GTEx donors (807 samples total) (Fig. 1A, B). For donor and sample information and regions targeted see (Datasets S1 & S2, respectively). The raw data have been deposited in a controlled-access public repository (dbGaP accession phs001746.v2.p1) linked to GTEx identifiers. We achieved high capture efficiency (*SI Appendix*, Fig. S1A, B, C); over 90% of targeted regions were covered at 30x sequencing depth in nearly all 807 samples (Fig. 1C, D, *SI Appendix*, Fig. S1B). Data on read counts, alignment efficiency, bisulfite conversion efficiency, and duplication rate are provided (Dataset S3). A small subset of difficult to capture regions failed to meet coverage criteria in all libraries (*SI Appendix*, Fig. S1C, Dataset S4). A set of Y-chromosome regions included in the capture enabled us to confirm that all 807 samples are of the correct sex (*SI Appendix*, Fig. S1D), indicating reliable sample handing.

**Fig. 1.**
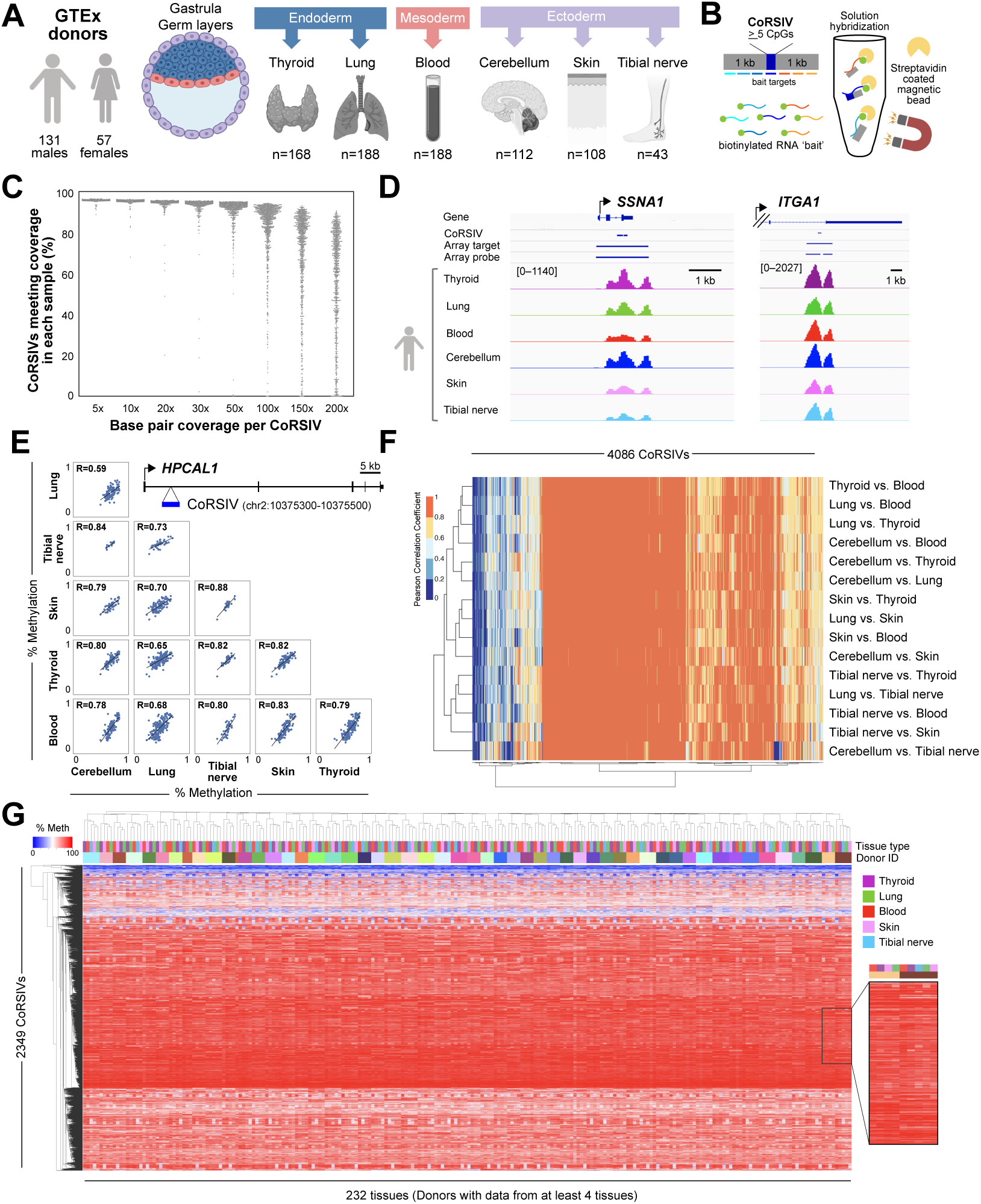
Target-capture bisulfite sequencing in 807 GTEx samples confirms systemic interindividual epigenetic variation at CoRSIVs. (**A)** DNA samples were obtained from multiple tissues (representing the three embryonic germ layers) from each of 188 GTEx donors. (**B)** CoRSIV capture process using Agilent reagents. (**C)** Percentage of CoRSIVs for which target-capture bisulfite sequencing achieved various read depths; each point represents one of 807 samples. (**D)** Plots of read depth at two target regions illustrate specificity of targeting across all six tissues.Y-axis scales are same for each region, and indicated for thyroid. (**E)** Scatter plots between all possible tissue pairs illustrate high inter-tissue correlations at a CoRSIV within *HPCAL1*. (**F)** Heat map of inter-tissue correlations across 4,086 CoRSIVs shows generally high correlation coefficients between all possible tissue pairs. (**G)** For the 232 tissue samples from 53 donors with data on at least 4 tissues (excluding cerebellum), unsupervised hierarchical clustering of methylation data at 2,349 fully informative CoRSIVs groups perfectly by donor.

CoRSIVs were identified based on unbiased genome-wide assessment of DNA methylation in thyroid, heart, and brain (37). Our first goal, therefore, was to examine additional tissues to confirm systemic interindividual variation (SIV) at these regions. High inter-tissue correlation in DNA methylation is the hallmark of SIV (Fig. 1E). Of the 4,641 genic CoRSIVs targeted, the 4,086 that satisfied coverage criteria in at least 10 donors in every possible pair of tissues were evaluated. Most of these showed high positive inter-tissue correlations (Pearson R>0.6) across all possible tissue pairs (Fig. 1F, *SI Appendix*, Fig. S1E, Dataset S5), confirming SIV. Accordingly, unsupervised clustering of methylation data at the 2,349 CoRSIVs covered in all 5 tissues (except cerebellum) across 53 donors grouped perfectly by the donor (Fig. 1G, Dataset S6). This clustering was not associated with sample-level variation in capture efficiency (Dataset S7). As DNA methylation in the cerebellum often differs from that in other brain regions (39), including cerebellum in this analysis resulted in a minor cerebellum cluster (*SI Appendix*, Fig. S1F); nonetheless, high inter-tissue correlations were maintained (*SI Appendix*, Fig. S1G). Of greatest relevance to epigenetic epidemiology, CoRSIV-specific scatter plots of methylation in brain, thyroid, skin, lung, and nerve versus blood show that methylation in blood generally serves as a proxy for methylation in other tissues (<u>five tissues vs. blood</u>). By comparison, in an HM450 study of 122 individuals (39), correlations between methylation in 4 brain regions vs. blood averaged only 0.2 and rarely exceeded 0.5. Although the inter-tissue scatter plots at CoRSIVs commonly show either a uniform distribution or three clusters (suggesting a single-genotype effect) (*SI Appendix*, Fig. S2), other patterns observed include 2, 4, and 5 distinct clusters (*SI Appendix*, Fig. S3). Consistent with our earlier study (37), in all six tissues almost every CoRSIV displayed an interindividual methylation range >20% (median range 40-42%) (*SI Appendix*, Fig. S4). Together, these results validate these CoRSIVs as systemic individual variants, essentially epigenetic polymorphisms.

### Gene expression in internal organs correlates with CoRSIV methylation in blood

Compared to genetic epidemiology, epigenetic epidemiology is complicated by the inherent tissue-specificity of epigenetic regulation (5). Because nearly all EWAS are based on measuring methylation in peripheral blood DNA, attempts to discover associations with, for example, Alzheimer’s disease (9) or schizophrenia (40) are implicitly predicated on the assumption that methylation variants in blood associate with epigenetic regulation in the brain. Of those on the Illumina arrays, however, such probes are the exception (39, 41). We therefore used our target capture bisulfite sequencing data and transcriptional profiling (RNA-seq) data from GTEx to test for cross-tissue correlations between CoRSIV methylation and expression of associated genes.

Of 3,768 CoRSIV-associated genes, over half showed appreciable expression in at least 5 of the six tissues under consideration (*SI Appendix*, Fig. S5A, B). Tibial nerve was excluded from this analysis due to low sample size; for each other tissue, both CoRSIV methylation and gene expression data were available for at least 60 individuals (*SI Appendix*, Fig. S5C). Tissues that are difficult to sample non-invasively (thyroid, lung, and cerebellum) were considered ‘target’ tissues. Within each of these we identified all CoRSIV-gene pairs for which gene expression is associated with CoRSIV methylation (FDR<0.05) (*SI Appendix*, Fig. S6A, B show two examples). Relative to those within a gene body, CoRSIVs located within 3 kb of either the 5’ or 3’ end of a gene showed predominantly negative correlations between methylation and gene expression (OR=2.84, P = 0.002) (*SI Appendix*, Fig. S6C).

For each CoRSIV-gene pair showing an expression vs. methylation association in a target tissue, we next asked whether methylation measured in easily accessible ‘surrogate’ tissues (blood or skin) is associated with expression in the target tissue. Of 156 genes for which expression was correlated with CoRSIV methylation in thyroid, for example, 122 (75%) showed a significant correlation and in the same direction when methylation in blood was used as the independent variable (*SI Appendix*, Fig. S6D). Likewise, in lung and cerebellum at least 75% of all methylation-expression correlations were detected when methylation in blood was used to infer expression (*SI Appendix*, Fig. S6D). In the other surrogate tissue, skin, this figure was slightly lower (60%). These data demonstrate that, at gene-associated CoRSIVs, methylation measurements in easily accessible tissues like blood can be used to draw inferences about epigenetic regulation in internal organs, a major advantage for epigenetic epidemiology.

### Genetic influences on methylation at genic CoRSIVs are strong and biased

The Genetics of DNA Methylation Consortium (GoDMC) recently analyzed HM450 and genotyping data on nearly 33,000 people in 36 cohorts (42) and documented mostly modest effects; for 75% of the *cis* mQTL associations the genetic variant explained less than 5% of the variance in methylation. In the largest unbiased study of human mQTL, Busche et al. (43) performed whole-genome bisulfite sequencing in 43 female twins and concluded environment, not genetics, is the main source of interindividual variation in DNA methylation.

We wondered to what extent individual variation in CoRSIV methylation is explained by genetic variation in *cis*. Within each CoRSIV, methylation of multiple CpGs is highly correlated (37); we therefore tested for genetic associations with average CoRSIV methylation, rather than at the CpG level. Also, given the multiplicity of mQTL associations at each CoRSIV (median 22 SNVs with P<10^-10^ per CoRSIV, *SI Appendix*, Fig. S7), rather than attempt to detect all possible SNV-CoRSIV associations we employed the Simes correction (44) to identify the single SNV most strongly associated with methylation at each CoRSIV (lowest p value, adjusted for multiple testing) (Fig. 2A, B, *SI Appendix*, Fig. S8, Dataset S8; listed p values are adjusted for multiple testing.) This approach conservatively tests each CoRSIV for evidence of genetic influence on its methylation, and is much more powerful than those we were able to employ in our earlier study (37) based on just 10 individuals.

**Fig. 2.**
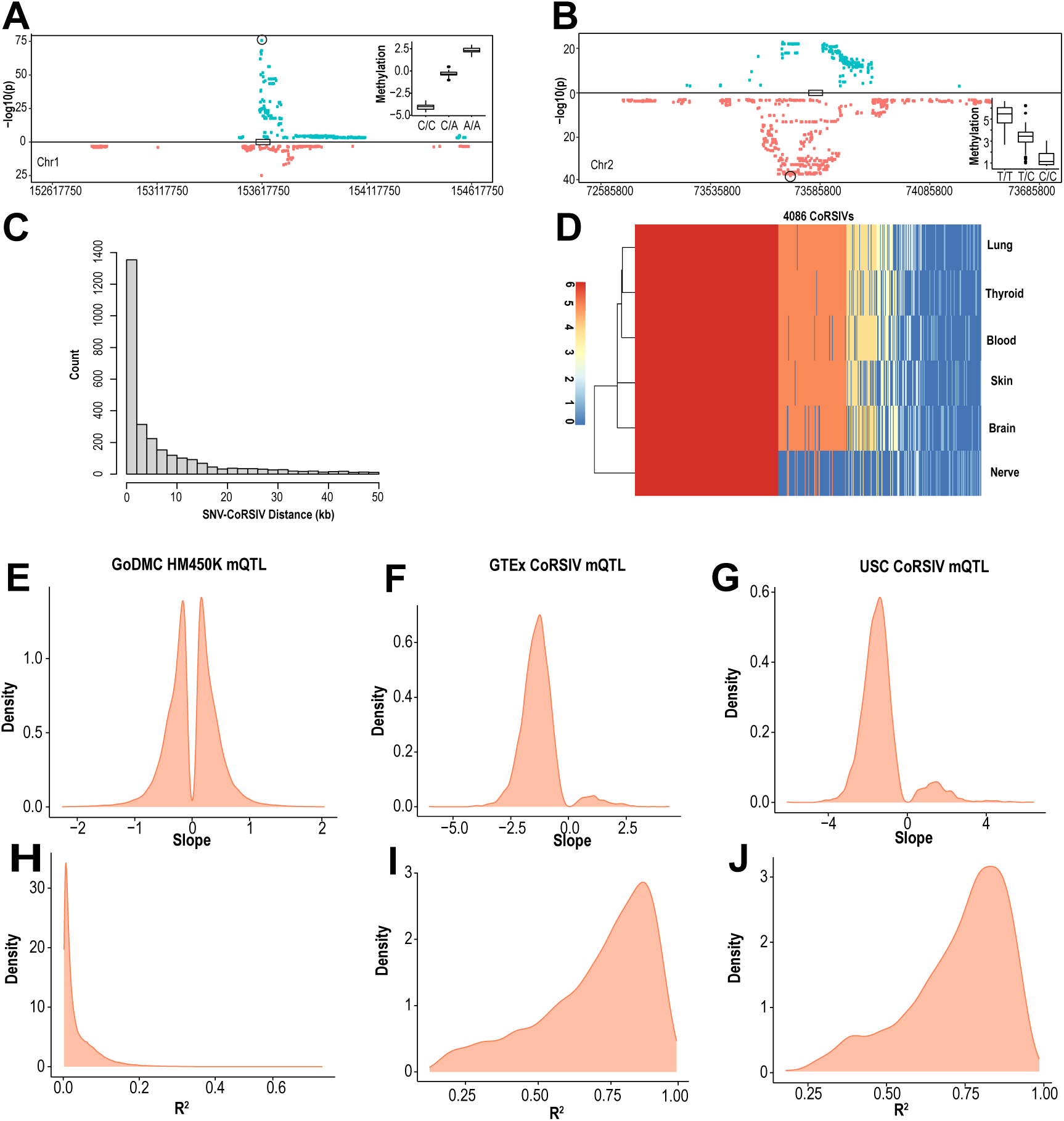
Genetic influences on CoRSIV methylation are strong and biased. (**A) (B)** Representative plots of mQTL associations at individual CoRSIVs on chromosomes 1 and 2, respectively. Significant associations are shown for all SNVs within 1Mb of each CoRSIV; positive and negative beta coefficients are plotted in blue and red, respectively. The most significant SNV (Simes SNV) is circled. Insets show average CoRSIV methylation vs. Simes SNV genotype. (**C)** Distribution of distances between CoRSIVs and corresponding Simes SNVs. **(D)** For each of 4,086 CoRSIVs, heat map depicts the number of tissues in which the Simes SNV falls within the same haplotype block, illustrating the largely systemic nature of mQTL at CoRSIVs. (E) Distribution of beta coefficients of significant Simes mQTL associations for the GoDMC blood mQTL data (42) (**F)** Distribution of beta coefficients of significant Simes mQTL associations at 3,723 CoRSIVs in blood DNA from 188 GTEx donors. **(G)** Distribution of beta coefficients of significant Simes mQTL associations across 2,939 CoRSIVs in blood DNA from 47 newborns (USC). **(H)** Distribution of Simes mQTL R^2^ (goodness of fit) for the GoDMC data. **(I)** Distribution of Simes mQTL R^2^ at CoRSIVs (GTEx, blood). **(J)** Distribution of Simes mQTL R^2^ at CoRSIVs (USC samples).

Although we tested all SNVs within 1 Mb, ‘Simes SNVs’ were generally proximal to the CoRSIV, 72% within 10 kb (Fig. 2C, *SI Appendix*, Fig. S9). Remarkably, although the Simes procedure was carried out independently in each tissue, at each CoRSIV the exact same SNV in many cases yielded the strongest mQTL association in all or most of the tissues (*SI Appendix*, Fig. S10A, B). When we asked how often the Simes SNV was within the same haplotype block in all or most tissues, concordance was even stronger (Fig. 2D), indicating the systemic nature of genetic influences on methylation at genic CoRSIVs.

Previous studies of mQTL using the HM450 array (22, 42) consistently report beta coefficients balanced on both sides of zero, as we found by employing the Simes procedure to the GoDMC data (Fig. 2E). Conversely, most *cis* mQTL associations at genic CoRSIVs show a negative beta coefficient (i.e., the major allele is associated with higher methylation) (Fig. 2F). This imbalance held not just for Simes SNVs, but for all mQTL SNVs (*SI Appendix*, Fig. S11). The strength of mQTL associations at genic CoRSIVs also appears to be without precedent (22, 42). In the GoDMC data, for example, few Simes mQTL associations show an R^2^ > 0.2 (Fig. 2H); at CoRSIVs, the median R^2^ = 0.76 (Fig. 2I, *SI Appendix*, Fig. S12). This tendency for high-R^2^ mQTL was largely independent of the distance between CoRSIV and SNV (*SI Appendix*, Fig. S13).

We made several attempts to disprove these surprising findings. Though unlikely (because each CoRSIV contains at least 5 CpGs (37)), we first asked whether the strong mQTL effects could be caused by SNVs abrogating CpG sites within CoRSIVs. Of SNVs present in our sample of 188 individuals, at least one did overlap a CpG within most of the CoRSIVs we surveyed. The distributions of beta coefficient and R^2^ values of Simes mQTL associations for the 1,155 CoRSIVs without any such overlaps, however, were nearly identical to those of the 2,759 with SNV-CpG overlaps (*SI Appendix*, Fig. S14). We next asked whether, instead of affecting CpG sites, SNVs within CoRSIVs might introduce an artifact by compromising the binding of the baits used for target capture. Despite their small size (median 200 bp), most CoRSIVs contain 2 or more SNVs (*SI Appendix*, Fig. S15A); however, neither the beta coefficients nor the R^2^ values of the Simes mQTL associations were strongly associated with the number of SNVs per CoRSIV (*SI Appendix*, Fig. S15B, C). Together, these data indicate that the strong and biased mQTL effects we detected are not due to SNVs within CoRSIVs.

For a complementary analysis, we employed a haplotype-based approach to assess genetic influences on CoRSIV methylation. We used phased genotype data from GTEx to infer each individual’s haplotype within the haplotype block overlapping each CoRSIV and assessed correlations between CoRSIV methylation and haplotype allele sum (sum of minor alleles in each individual) (*SI Appendix*, Fig. S16A). This analysis yielded a preponderance of negative coefficients, and local haplotype explained much of the variance in methylation (median R^2^ = 0.43) (*SI Appendix*, Fig. S16B, Dataset S9), consistent with the mQTL analysis.

Lastly, to independently validate genetic effects on CoRSIV methylation we performed CoRSIV-capture bisulfite-sequencing and SNV genotyping in 47 individuals from a different (non-GTEx) population (USC cohort). To ensure computational independence, a separate member of our laboratory wrote new code for the Simes mQTL analysis. The USC results corroborated the negative bias and high R^2^ of mQTL effects at CoRSIVs (Fig. 2G, J). An independently performed haplotype-based analysis likewise corroborated the results obtained on the GTEx samples (*SI Appendix*, Fig. S16C). Together, these additional analyses and data indicate that the strong and biased genetic influences on methylation at these CoRSIVs are genuine.

We wondered how the total amount of mQTL we detected at genic CoRSIVs compares with that reported by the GoDMC (42), which used HM450 arrays to study 33,000 people. With 3 genotype calls possible at each SNV, the average methylation difference (delta) associated with each SNV can be calculated from the mQTL beta coefficient (*SI Appendix*, Fig. S17A). And, since the mQTL R^2^ measures what proportion of this delta is explained by SNV genotype, the product (delta)x(R^2^) measures the absolute methylation variation explained by SNV genotype. To make our results interpretable, we initially assessed (delta)x(R^2^) based on beta values (rather than using the M-value transformation). Across all CoRSIV mQTLs (P < 10^-10^), median (delta)x(R^2^) was 24.6% methylation (*SI Appendix*, Fig. S17B); for a CoRSIV with an R^2^ near the median (0.76), this equates to an interindividual range of 32.4% methylation, within the normal range for CoRSIVs (*SI Appendix*, Fig. S4). To compare our results with those of GoDMC (42), whose coefficients were provided based on M values, we repeated our analysis after applying the M value transformation. At the CoRSIVs we assayed, the total methylation variance explained by genetics (sum of (delta)x(R^2^)) was 72-fold greater than that detected by GoDMC (42) (*SI Appendix*, Fig. S17C, D, E), the largest study of human mQTL ever reported.

Genetic influences on tissue-specific expression (eQTL) can be mediated by mQTL (23, 45). Given the strong mQTL effects at genic CoRSIVs, we used data from GTEx (46) to ask whether Simes SNVs are enriched for eQTL. Consistent with the analysis of GTEx data overall (46) many eQTL effects were shared among non-brain tissues, whereas eQTL associations in brain and blood were more distinct (*SI Appendix*, Fig. S18A). Relative to all common variants, which have a 50% chance of being associated with expression of a nearby gene (46), a bootstrapping analysis indicated that Simes SNVs are 3.4-fold more likely to show eQTL effects (*SI Appendix*, Fig. S18B). The distributions of magnitude, slope, and SNV-eGene distance for eQTL effects at Simes SNVs were similar to those of all common variants (*SI Appendix*, Fig. S18C, D). Future studies will be required to determine if the enriched eQTL at Simes SNVs is in some cases mediated by CoRSIV mQTL.

### CoRSIVs occur in genomic regions with far-reaching enrichments in transposable elements

The earliest known examples of systemic interindividual epigenetic variants in mammals are mouse metastable epialleles such as *agouti viable yellow* and *axin fused*, both of which resulted from retrotransposition of an intracisternal-A particle (an LTR-retrotransposon) (47, 48). We previously showed that CoRSIVs are enriched for direct overlaps with LINE, SINE, and ERV retrotransposons (37); we provide a more granular analysis of those overlaps here (*SI Appendix*, Fig. S19). Given the ability of transposable elements for long-range regulation of transcriptional and epigenetic dynamics in the early embryo (49, 50) we asked whether the exceptional behavior of CoRSIVs might be associated with specific classes of repetitive elements working over long genomic distances.

Relative to a set of control regions matched to genic CoRSIVs by chromosome, size, and CpG density (37), in regions flanking genic CoRSIVs we detected long-range depletion of CpG islands and enrichments of specific classes of LINE and LTR retrotransposons (Fig. 3A, Dataset S10). Similar and stronger enrichments were detected in comparison with size-matched tissue-differentially methylated regions (tDMRs) (37) (*SI Appendix*, Fig. S20). Interestingly, enrichments relative to control regions (Fig. 3A) were strongest among the evolutionarily youngest subclasses, the LINE1-PA elements (51) among LINEs, and ERV-K elements (50) among LTRs.

**Fig. 3.**
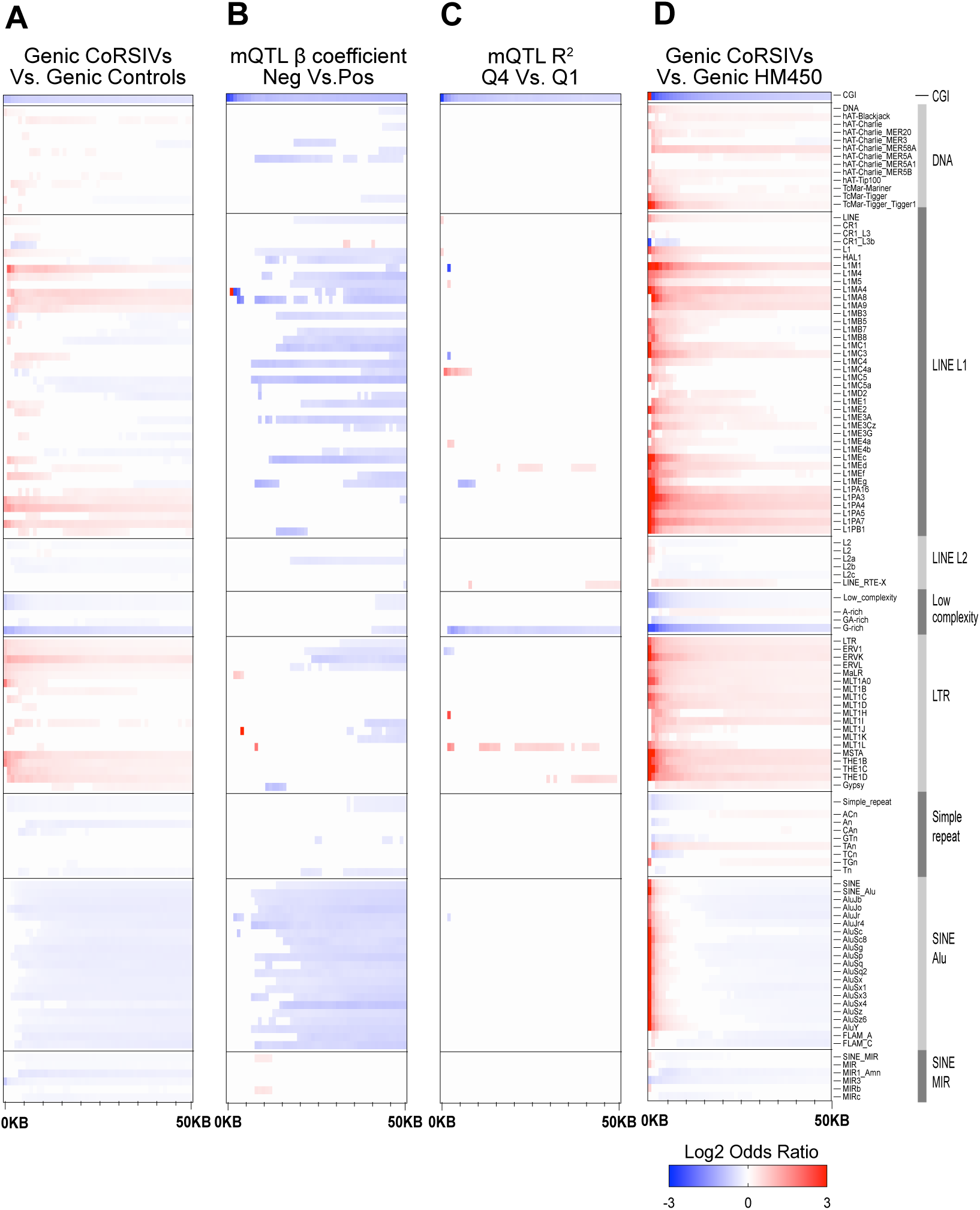
Genic CoRSIV-flanking regions show long-range enrichments and depletions for specific classes of transposable elements. **(A)** Using 1 Kb step sizes, each plot shows significant enrichments or depletions for CpG islands (CGI) and subclasses within each of 8 classes of transposable element within 50 Kb of genic CoRSIVs. Compared to control regions, CoRSIV-flanking regions show long range depletion of CpG islands and enrichment of specific classes of LINEs and LTRs. **(B)** Compared to CoRSIVs showing a positive mQTL beta coefficient, those with negative coefficients are depleted for CpG islands and show long-range depletion of specific LINE1s and all subclasses of Alus. **(C)** The strength of mQTL associations at CoRSIVs (R^2^ in 4^th^ vs. 1^st^ quartile) is not associated with widespread differences in genomic content of transposable elements. **(D)** Compared to regions in which HM450 probes are located, CoRSIVs show short- and long-range enrichments for many subclasses of LINE1 and LTR retrotransposons.

We next asked whether either the negative bias (i.e., the major allele associating with higher methylation) or the strength of mQTL associations at CoRSIVs might be associated with transposable elements in flanking genomic regions. Compared to genic CoRSIVs showing a positive mQTL beta coefficient, those characterized by negative coefficients were depleted for CpG islands (Fig. 3B). There were no robust short-range associations of transposable elements with ‘negative mQTL’ CoRSIVs; rather, at distances > 5-10kb from the origin they show extensive long-range depletion of specific LINE1 and all classes of Alu elements (Fig. 3B, Dataset S11). Surprisingly, the strength of mQTL at genic CoRSIVs was not associated with widespread differences in genomic content of transposable elements. Relative to those in the bottom quartile for R^2^, mQTL effects in the top quartile showed proximal and long-range depletion in just CpG islands and G-rich low-complexity repeats (Fig. 3C, Dataset S12).

As most human mQTL data are based on the HM450 array, we next compared genomic regions flanking genic CoRSIVs with those flanking genic HM450 probes, finding striking differences. Although the HM450 array specifically targets CpG islands, these are more strongly enriched within 1 kb of genic CoRSIVs (Fig. 3D, Dataset S13); at greater distances, CoRSIV-flanking regions are relatively depleted of CpG islands. Compared to genomic regions containing genic HM450 probes, those housing genic CoRSIVs show strong short-range (1-2 kb) enrichments in LINE1, LTR, and Alu elements (Fig. 3D). The LINE1 and LTR enrichments gradually weaken but extend to at least 50 kb from the origin. Enrichments for Alu extend only to ∼5 kb; at greater distances, regions flanking genic CoRSIVs are relatively depleted (Fig. 3D). These enrichments were not unique to genic CoRSIVs; the full set of 9,926 CoRSIVs showed similar patterns of enrichment relative to matched control regions, tDMRs, and HM450 probes (*SI Appendix*, Fig. S21). These observations suggest a straightforward explanation for the strong and biased mQTL effects at CoRSIVs. To limit hybridization artifacts, the Illumina methylation arrays avoided genomic regions rich in transposable elements. But these are the same regions in which SIV tends to occur. Given the potentially deleterious consequences of transcriptional activation of retrotransposons, the strong and negative mQTL beta coefficients at CoRSIVs could reflect evolutionary selection for genetic variants favoring their methylation and silencing. In support of this, values of Tajima’s D (a test statistic assessing evidence of evolutionary selection) are higher in CoRSIVs compared to control, tDMR, or HM450 probe regions (*SI Appendix*, Fig. S22, Dataset S14).

### CoRSIV flanking regions are enriched for heritability of disease

Across diverse outcomes including Alzheimer’s (23), chronic obstructive pulmonary disease (52), obsessive-compulsive disorder (53), and cardiovascular disease (54), integrative analyses of GWAS and DNA methylation profiling data increasingly indicate that mQTL mediates associations between genetic variation and risk of disease. We therefore asked whether the strong mQTL effects identified at genic CoRSIVs are associated with genetic variants identified by GWAS. Indeed, permutation testing indicates that SNVs identified in our mQTL analysis are enriched for SNVs implicated in metabolic, hematological, anthropometric, cardiovascular, immune, neurological, and various other diseases (Fig. 4 A, B, Dataset S15). By contrast, despite an abundance of CoRSIV-associated genes linked to cancer (37), no enrichment was found relative to cancer GWAS SNVs (Fig. 4 A, B). Notably, a recent analysis employing these same categories (24) found nearly opposite categorical enrichments with *trans*-mQTL loci. With the caveat that 90% of GWAS alleles impact multiple traits (55), it is interesting that cancer traits are not enriched. This may indicate that CoRSIV methylation plays no role in this maladaptive phenotype, or reflect dilution of effects across multiple cancer subtypes and various genetic pathways leading to cancer (56). Overall, and particularly considering that Simes SNVs are enriched for eQTL, these results are consistent with the possibility that human genetic variants influence disease risk via mQTL effects at CoRSIVs.

**Fig. 4.**
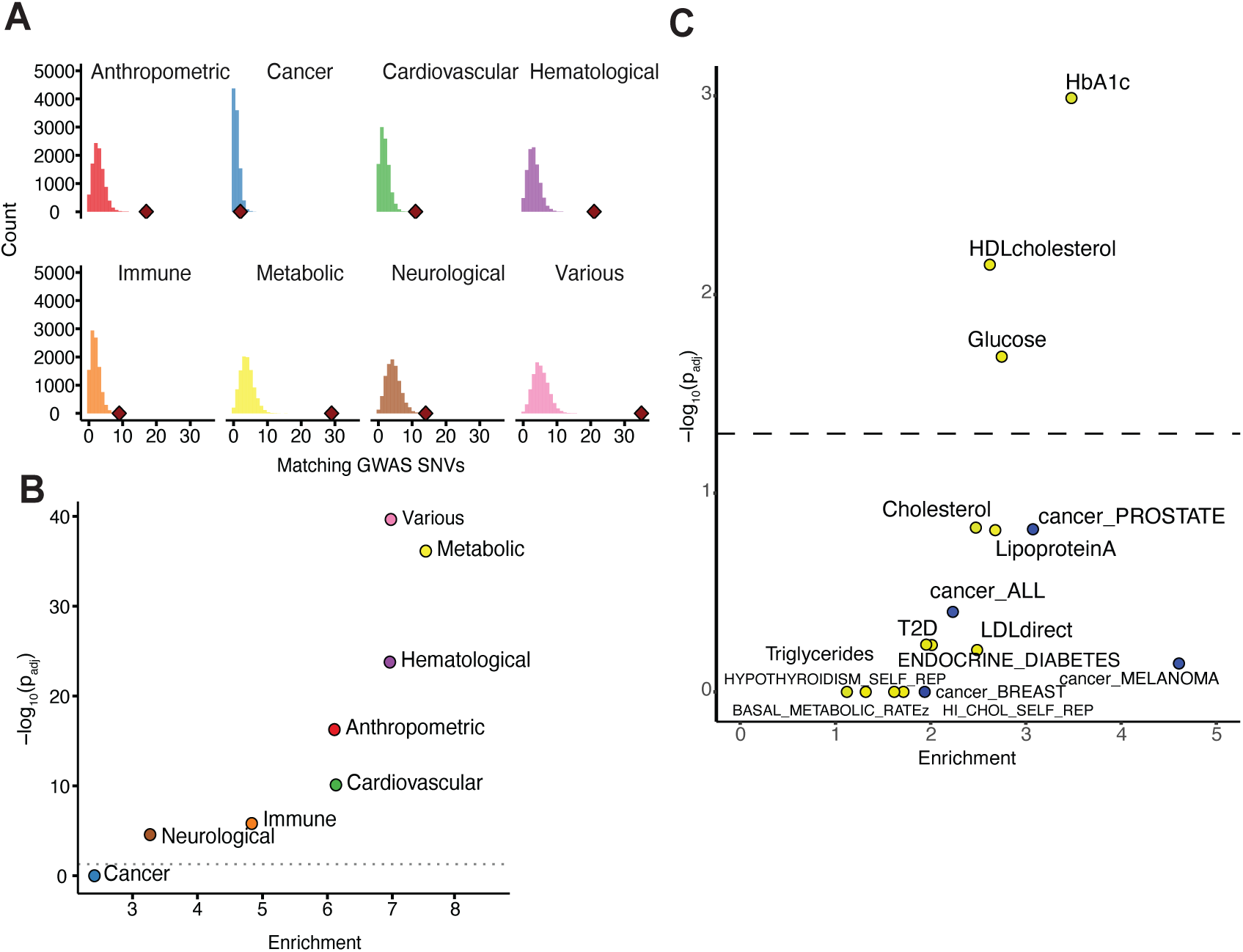
CoRSIV mQTL SNVs are enriched for GWAS associations. **(A)** Within each of 8 disease/phenotype categories, the histogram shows the null distribution obtained by permutation testing for overlap of GWAS SNVs with SNVs randomly sampled within 1Mb of each CoRSIV. The red diamond shows the actual number of overlaps between CoRSIV mQTL SNVs and GWAS SNVs. Numbers of GWAS SNVs considered in each category are anthropometric: 8106, cancer: 3,163, cardiovascular: 4,816, hematological: 7,461, immune: 5,263, metabolic: 10,121, neurological: 14,741, and various: 14,573. **(B)** Statistical significance (Bonferroni-adjusted p-value) vs. fold enrichments for the analysis in (A). Strong and statistically significant enrichments were found for all outcomes except cancer. **(C)** Statistical significance (Bonferroni-adjusted p-value) vs. fold enrichments for 8 metabolic traits and 4 cancer outcomes from the LDSC analysis confirms that the vicinity of CoRSIVs is enriched for heritability of metabolic traits.

As a complementary analysis, we used LD score regression (LDSC) (57) to determine if, in the vicinity of genic CoRSIVs, there is enrichment for heritability of metabolic phenotypes and cancer. GWAS summary statistics from the UK Biobank representing 12 metabolic traits and 4 cancer outcomes were downloaded (58). As nearly all Simes SNVs are within 20 kb of their associated CoRSIV (Fig. 2C), we evaluated genomic regions encompassing genic CoRSIVs +/- 20 kb. Consistent with our results based on direct overlap with Simes SNVs, individual LDSC models focused on each outcome detected significant enrichment for 3 metabolic outcomes (HbA1c, HDL cholesterol, and glucose) but none for cancer (Fig. 4C). As suggested by Finucane et al (57), we repeated these analyses including in each a full ‘baseline’ model comprising 53 sequence and epigenomic features. Enrichment for heritability of two of the metabolic traits, HbA1c and HDL cholesterol, was attenuated but remained significant (*SI Appendix*, Fig. S23A). The baseline-adjusted analysis (*SI Appendix*, Fig. S23B) confirmed strong evolutionary conservation in the vicinity of genic CoRSIVs. Also, significant enrichments for coding regions and transcription start sites may explain the attenuated associations with metabolic outcomes. Regardless, we would argue that because CoRSIVs were identified based solely on SIV in DNA methylation it is inappropriate to penalize them for association with genic and regulatory features. Hence, the LDSC results corroborate that CoRSIV-flanking regions are enriched for heritability of metabolic disease.

## Discussion

Following up on our previous screen for human CoRSIVs (37) here we have, for the first time, demonstrated the feasibility of studying these regions at the population level using target-capture bisulfite sequencing. Performing these analyses on donors from GTEx allowed us to integrate our methylation data with genome sequence and gene expression data on these same individuals. As expected, our results validated SIV at the CoRSIVs we analyzed, and indicate the ability to use methylation profiling in peripheral blood to draw inferences about epigenetic regulation in various organs of the body. More surprisingly, our analyses of genetic influences on CoRSIV methylation indicate an unprecedented level of mQTL at these regions. Also unlike previous reports, our mQTL analysis showed strongly biased beta coefficients (i.e., the major allele associated with higher methylation). Lastly, we found evidence that genomic regions encompassing CoRSIVs are enriched for the heritability of human disease traits.

Though unprecedented, the extremely strong mQTL effects at the CoRSIVs we surveyed are unsurprising. Because variation at each SNV is fixed (ranging from 0 – 2 copies of the minor allele), the best way to increase the power of mQTL detection is to focus on CpG sites with the greatest interindividual range of DNA methylation. Other than our work (37, 59, 60), we are not aware of previous studies that took this approach. Instead, nearly all investigations of human mQTL have employed Illumina arrays (22), which do not target interindividual variants. One may question the validity of quantitatively comparing our mQTL results with those of GoDMC (42). After all, GoDMC analyzed HM450 data on 420,000 CpG sites across nearly 33,000 individuals, whereas we analyzed target-capture bisulfite sequencing data on 4,086 CoRSIVs in just 188 individuals. But although the targeted regions and studied populations differ, both analyses employed the same statistical method for mQTL detection. Because GoDMC performed their mQTL analyses using M values (a transformation of the Beta value intended to improve normality), we also transformed our percent methylation data to M values for this comparison. Therefore, despite the different approaches and vastly dissimilar numbers of subjects considered, our analysis is quantitatively comparable to that of Min et al. (42). Our ability to detect more mQTL than ever before despite surveying a much smaller number of CpG sites than on the Illumina arrays speaks to the importance of targeting the right CpGs. Known human CoRSIVs comprise just 0.1% of the genome; whilst some may question the wisdom of focusing on such a small fraction of genomic CpG sites, common human sequence variants comprise only ∼0.3% of the genome (26) but have been a major focus of the GWAS field for the last 20 years.

In addition to the extremely strong mQTL effects at genic CoRSIVs, we are not aware of previous studies showing a bias in mQTL regression coefficients (Fig. 2, F & G). The mQTL bias at genic CoRSIVs reflects that the major allele is generally associated with higher methylation. This is consistent with the enrichment of L1 and LTR transposable elements in the vicinity of CoRSIVs (Fig. 3), because these tend to locate in heterochromatic regions (61). During human pre-implantation development, when methylation at CoRSIVs is thought to be established (37, 62), widespread genomic de-methylation leads to transient transcriptional activation of transposable elements, prior to their re-methylation and silencing in differentiated tissues (63). The high density of L1 and LTR retrotransposons in CoRSIV-flanking regions therefore raises the question of whether mQTL effects at CoRSIVs reflect modulation of the *establishment* of de novo or early embryonic *maintenance* of existing zygotic methylation. In this regard, it is striking that, in mice, L1 elements and IAPs (a class of LTR retrotransposons) are preferentially methylated in sperm and not oocytes, whereas Alus show the opposite pattern (methylated in oocytes but not in sperm) (64). These observations mirror our data on transposable element enrichments in regions flanking CoRSIVs (Fig. 3A). The biased mQTL beta coefficients at CoRSIVs lead us to speculate that they could reflect evolutionary selection for genetic variants that maintain methylation marks in the paternal genome, potentiating transgenerational epigenetic inheritance as observed at the murine metastable epiallele *axin fused* (65).

As DNA methylation can act as an intermediary molecular mechanism linking genetic variation to tissue-specific transcriptional regulation (23, 45), mQTLs may provide mechanistic insights into how genetic variants influence gene expression. In this regard, the dramatically different nature of mQTL effects at genic CoRSIVs, in terms of both strength and allelic bias, indicates that we have uncovered a fundamentally different component of epigenetic regulation compared to CpGs represented on the HM450 and EPIC arrays, which have largely been the focus of the field (22). Also, our observation that SNVs wielding the strongest mQTL effects at genic CoRSIVs are enriched for eQTL suggests a mechanistic pathway in which genetic effects on CoRSIV methylation modulate tissue-specific gene expression. On the other hand, 16% of CoRSIVs showed weak effects explaining less than half of the interindividual variation (Fig. 2I). These are candidate metastable epialleles. Future large human studies can better characterize genetic effects on CoRSIV methylation and elucidate true epipolymorphisms (i.e. metastable epialleles) at which a majority of interindividual epigenetic variation is unexplained by genetics, such as the non-coding RNA *nc886* (also known as *VTRNA2-1*) (17, 66). Combining data on such regions with those on recently identified murine metastable epialleles (67) may enable comparative genomic approaches to characterize sequence features that confer epigenetic metastability, informing *in silico* identification of metastable epialleles in other mammalian species.

Many important questions remain unanswered by our study. Our initial identification of CoRSIVs was based on ten Caucasian individuals. Reflecting the GTEx study overall, 90% of the donors included in this current study are also Caucasian. Although our previous studies (37, 59, 60) indicate that SIV regions identified in Caucasians generally also show SIV in other ethnic groups, future studies screening for SIV directly in non-Caucasian populations may identify CoRSIVs specific to other ethnic groups. Also consistent with the GTEx study population overall, most donors studied here were between 50-70 years old (Dataset S1). Considering the influence of age on epigenetic marks (12), one might ask to what extent interindividual variation at CoRSIVs is influenced by age. Notably, the validation studies we performed to corroborate mQTL effects at CoRSIVs (Fig. 2, G & J) were based on peripheral blood of newborns yet showed nearly identical profiles of mQTL slope and variance explained, arguing that age is not a major factor in the regulation of systemic interindividual epigenetic variation. Compared to our initial screen which surveyed thyroid, heart, and cerebellum, here we evaluated SIV in 4 additional tissues, with at least one representing each germ layer lineage (Fig. 1A). Hence, whereas our results confirm high inter-tissue correlation coefficients across most tissue pairs for ∼90% of genic CoRSIVs (Fig. 1F) many more tissues and cell types remain to be evaluated. The small fraction of genic CoRSIVs with low inter-tissue correlations (Fig. 1F) may reflect false positives in our original screen, or possibly exhibit interindividual variation across specific tissue lineages not evaluated here.

The generally strong mQTL at CoRSIVs is not due to the systemic nature of their interindividual variation. Most of these same regions would have been detected if, instead of our original three-tissue screen (37) we had conducted an unbiased genome-wide screen for interindividual variation in, say, peripheral blood leukocytes. In addition to CoRSIVs, such an experiment would detect interindividual variants specific to blood. Rather than interindividual variation intrinsic to leukocytes, however, many of these reflect interindividual variation in leukocyte composition (ratio of B cells to T cells, for example) (68). We would argue that such variants are not *bona fide* interindividual epigenetic variants. Because most human tissues exhibit such cellular heterogeneity, the specific composition of which can differ among individuals and disease states, interindividual variation observed in just one tissue is difficult to interpret. CoRSIVs, on the other hand, are unaffected by individual differences in tissue cellular composition (37); like sequence variants, they are stable epigenetic variants intrinsic to essentially all cells in an individual. The CpG methylation profile at CoRSIVs can therefore reasonably be considered a readout of an individual’s epigenome, enabling adoption of concepts and applications developed for genomics, such as GWAS. Given the strong influence of genetics on methylation at CoRSIVs, one might ask whether profiling CoRSIV methylation offers additional information beyond that obtained by genotyping. We anticipate many advantages. First, as multiple genetic variants influence methylation at each CoRSIV (*SI Appendix*, Fig. S7), CoRSIV methylation can be viewed as an integrative readout of these influences. Also, GWAS variants may logically be prioritized based on known mQTL effects at CoRSIVs, just as investigators now prioritize GWAS hits based on evidence of eQTL (69). In fact, mQTL effects at CoRSIVs may in some cases mediate eQTL. Lastly, whereas our current data on CoRSIV mQTL is based on a mostly Caucasian cohort in the US, it is possible that additional sources of variation (for example, due to periconceptional environment (37, 59, 60)) will be uncovered as CoRSIVs are studied in a broader range of ethnic and cultural contexts, providing insights into gene by environment interactions.

For over ten years the Illumina methylation platform has been the predominant tool for population studies of DNA methylation (22, 30). A major reason is that it interrogates a stable subset of CpG sites within the human genome (yielding one quantitative value for each), simplifying data sharing and integration across multiple studies and populations. Nonetheless, the platform has a major and undeniable shortcoming in the context of population epigenetics: most CpGs included do not show appreciable interindividual variation (33–36). Here we have shown that focusing on systemic methylation variants enables the identification of far stronger mQTL than has been detected by the Illumina arrays (42). We anticipate that the greater population variance at CoRSIVs will also improve the power of studies aiming to associate epigenetic variation with risk of disease. Generating the data to explore associations between CoRSIV methylation and a wide range of human diseases is beyond the scope of this study. However, though grossly underrepresented on the HM450 and EPIC arrays, CoRSIVs are often among top ‘hits’ in existing EWAS (70). Indeed, these stable (36, 60, 71), systemic epigenetic variants are already showing great promise for disease prediction (72–78). We suggest that improving the coverage of CoRSIVs would enhance the utility of the Illumina EPIC array for the study of population epigenetics. Additionally, we wish to make our validated human CoRSIV-capture reagents available to the field to facilitate the study of these systemic variants. The list of known human CoRSIVs is currently incomplete, and screening is underway to identify more, including in various ethnic groups.

## Materials and Methods

### Study samples

We obtained de-identified genomic DNA from multiple tissues of 188 donors in collaboration with NIH Genotype-Tissue Expression (GTEx) program (38) (total of 807 samples). Informed consent was obtained by GTEx, including authorization to release the patient’s medical records and social history, sequencing of the donor’s genome, and blanket consent for all future research using the donated tissue and resultant data. The donor and tissue information are available in Dataset S1 in the Supplementary Appendix. For the independent mQTL validation, newborn blood spots from pediatric glioblastoma cases and controls (47 samples total) were obtained from the California Biobank, using information from the California Cancer and Vital Statistics registries. Genotype data for the 188 individuals were generated by GTEx, and for the other 47 samples DNA extraction, preprocessing and genotyping were performed as previously described (79) (see the methods in supplementary appendix for more details).

### Target capture bisulfite sequencing and data processing

Out of 9,926 CoRSIVs previously reported (37), we included only those within 3000 base pairs from the body of a gene present in the PubTator (80) compendium, using BEDTOOLS (81) software, yielding 4641 CoRSIVs as targets for capture. The goal of using PubTator was to focus not just on known genes but on those most likely to be associated with a measurable phenotypic outcome. Libraries were made using the Agilent SureSelect Methyl-seq library kit with modifications (Design ID: S3163502). Capture design details and version history are available in the SI appendix, Materials and Methods. As for the data processing Bisulfite-sequencing reads were trimmed using Trim Galore, then mapped to the human genome build UCSC hg38 using the Bismark aligner (82). Uniquely mapped reads were retained for further analysis (see the methods in the supplementary appendix for more details).

### Evaluating genetic influences on CoRSIV methylation

Analysis of associations between CoRSIV DNA methylation and genetic variation in cis was performed relying on the Simes correction as described previously (44). Using the EMatrixQTL R package (83), Spearman rank correlation was computed for all SNVs within 1Mb of each CoRSIV, and the Simes correction was applied. Simes adjusted p-values for each CoRSIV were collected, and the false discovery rate (FDR) correction was applied across all CoRSIVs analyzed in each tissue, with significance achieved at FDR-adjusted p<0.05. To compare the summed total of mQTL detected at CoRSIVs vs. that reported by GoDMC (42), mQTL associations were identified with P < 10^-10^. This conservative P value was selected to avoid false positives, given the relatively small number of individuals in the GTEx CoRSIV analysis. To further evaluate genetic influence on CoRSIV methylation we used a haplotype-based approach. Phased genotype data from GTEx were used to infer each individual’s haplotype within the haplotype block overlapping each CoRSIV and assessed correlations between CoRSIV methylation and haplotype allele sum (see the methods in the supplementary appendix for more details).

### Data availability

The raw target capture bisulfite sequencing data for the 807 GTEx tissues (188 individuals) have been deposited to the AnVIL repository. Controlled access is administered through dbGaP (accession phs001746.v2.p1). The samples used in the mQTL validation analysis (USC cohort) are biospecimens from the California Biobank Program. Any uploading of genomic data and/or sharing of these biospecimens or individual data derived from these biospecimens would violate the statutory scheme of the California Health and Safety Code Sections 124980(j), 124991(b), (g), (h), and 103850 (a) and (d), which protect the confidential nature of biospecimens and individual data derived from biospecimens. Certain aggregate results from the USC cohort may be available from the authors by request.

## Acknowledgments

Funding for this project was provided by NIH/NIDDK (1R01DK111522), the Cancer Prevention and Research Institute of Texas (RP170295), and the USDA/ARS (CRIS 3092-5-001-059). The Functional Genomics core at Baylor College of Medicine, where the target-capture sequencing was done, is partially supported by NIH shared Instrument grant S10OD023469. The data used for the analyses described in this manuscript were obtained from the GTEx Portal on 03/01/2019 and/or dbGaP accession number phs000424.v8.p2. The GTEx Project was supported by the Common Fund of the Office of the Director of the National Institutes of Health, and by NCI, NHGRI, NHLBI, NIDA, NIMH, and NINDS. We are grateful that anonymous reviewers for two journals shared their time and expertise to help us improve the manuscript.

## Other Supplementary Materials for this manuscript include the following

Other Supplementary Materials for this manuscript include the following:
|  |  |
| --- | --- |
| Dataset 1 | GTEx Donor Information and Tissue Types |
| Dataset 2 | CoRSIVs targeted for Bisulfite Capture Sequencing (hg38) |
| Dataset 3 | CoRSIV Capture Sequencing Data QC Metrics |
| Dataset 4 | CoRSIVs which failed to meet coverage criteria in all libraries (hg38) |
| Dataset 5 | Inter-tissue Pearson correlation coefficients across six tissues (see Fig. 3F) |
| Dataset 6 | CoRSIV average methylation data for those adequately covered in all six tissues (see Fig. 1G) |
| Dataset 7 | Capture efficiency data do not associate with Fig. 1G clustering |
| Dataset 8 | Simes CoRSIV-SNV mQTL in GTEx Data (sorted by R-Squared) |
| Dataset 9 | Pearson correlation coefficients for haplotype allele sum vs. CoRSIV DNA methylation |
| Dataset 10 | Enrichment of repeat elements in Genic CoRSIVs vs. Controls |
| Dataset 11 | Enrichment of repeat elements in Genic CoRSIV mQTL slope Neg. Vs. Pos. |
| Dataset 12 | Enrichment of repeat elements in R-squared Q4 CoRSIV mQTLs. vs. Q1 CoRSIV mQTLs (Genic) |
| Dataset 13 | Enrichment of repeat elements in Genic CoRSIVs Vs. HM450K |
| Dataset 14 | Tajima's D Score comparison between CoRSIVs, Controls, tDMRs, HM450k |
| Dataset 15 | CoRSIV mQTL SNV association with GWAS SNVs |

## Materials and Methods

### CoRSIV Capture Design Versions

| **<u>V.</u>** | **<u>Agilent Design ID</u>** | **<u>Design Date</u>** | **<u>Included Regions</u>** | **<u>Agilent Design Selections</u>** | **<u>Size (Mbp)</u>** |
| --- | --- | --- | --- | --- | --- |
| 1.0 | S3163502 | 23-Aug-2018 | 4641 CoRSIVs<br>Sex specific Chr Y regions | 1x Tiling Density<br>Least Stringent Masking<br>Balanced Boosting | 9.045 |
| 2.0 | S3223244 | 16-Jul-2019 | 9926 CoRSIVs<br>SIV (1, 2)<br>ESS (2)<br>Sex-specific Chr Y regions<br>imprinting control regions (3) | 1x Tiling Density<br>Least Stringent Masking<br><u>Balanced Boosting</u> | 19.843 |
| 3.0 | S3295946 | 06-Aug-2020 | 9926 CoRSIVs<br>SIV(1, 2)<br>ESS (2)<br>Cell Composition estimation probes (4, 5)<br>Sex-specific Chr Y regions, imprinting control regions(3) | 1x Tiling Density<br>Least Stringent Masking<br>Optimized Performance<br>XT/XT2 boosting | 21.958 |
| 3.1 | S3332366 | 23-Feb-2021 | 9926 CoRSIVs<br>SIV(1, 2)<br>ESS (2)<br>Cell Composition estimation probes (4, 5)<br>Sex-specific Chr Y regions, imprinting control regions(3) | 1x Tiling Density<br>Least Stringent Masking<br>Optimized Performance<br>XT/XT2 boosting<br>Overnight Hybridization<br>Deleted ineffective baits from v.3.0. | 21.130 |

### Design of CoRSIV-capture reagent

Of the 9,926 CoRSIVs previously reported (*6*), to ensure adequate targeting we filtered to include only those within 3,000 base pairs (bp) from the body of a gene present in the Pubtator (*7*) compendium, using BEDTOOLS (*8*) software, yielding 4,641 CoRSIVs as targets for capture (Supplementary Table). For quality control purposes we included 10 regions on the Y chromosome to confirm the accurate biological sex of each sample. At each of the 4,641 CoRSIVs, the target region included flanking regions of 1,000 bp in each direction. We used the Agilent SureSelect online system to design a custom capture reagent, using the following options: balanced boosting, 1x tiling, and least stringent masking. Overall, our CoRSIV capture reagent (**Agilent Design ID: S3163502**) targeted 9.045 MB of the human genome (Supplementary Table 2), using 85,538 probes.

### Library preparation, capture, and sequencing

Individual libraries were made using the Agilent SureSelect Methyl-seq library kit, with modification. In brief, 1ug of genomic DNA was subject to shearing to 150-200bp in size using a Covaris sonicator. After purification through AMpure XP beads, end repair and A-Tailing was carried out. Then, 5ul of 15uM methylated library adaptor (IDT) was ligated to each sample, and the product with a size of 250-450bp was selected through Ampure XP beads.

Twelve libraries were pooled in equal proportions for target enrichment following an Agilent protocol (Sureselect Methyl-seq target enrichment system for Illumina multiplexed sequencing). After hybridization with probes (Agilent SureSelect, custom design), Dynabeads MyOne streptavidin T1 beads were used to bind the library. After several round of washes, the bound DNA was eluted in 0.1N NaOH and subjected to Bisulfite treatment using the EZ DNA Methylation Gold kit (Zymo Research). Final library was generated by amplification using Sureselect Methyl-seq PCR Master Mix and P5, P7 primers (Illumina). Sequencing was performed using an Illumina Novaseq 6000 at the Functional Genomics core, Department of Molecular and Human Genetics, Baylor College of Medicine.

### Data processing

Bisulfite-sequencing reads were trimmed using Trim Galore, then mapped to the human genome build UCSC hg38 using the Bismark aligner (*9*). Uniquely mapped reads were retained for further analysis. Duplicate reads were not removed, as recommended for capture experiments by the Bismark manual. CpG-level methylation was quantified using the Bismark pipeline. For each sample, average proportional DNA methylation was computed at each CoRSIV for which at least half of the CpGs were covered by at least 5x reads.

### Quality control assessment

To determine the proportion of ‘on-target’ reads, only those that mapped completely within a target region were counted; capture efficiency was calculated as the fraction of on-target reads divided by all uniquely mapped reads. To confirm the accuracy of the biological sex of each sample, coverage of chromosome Y control regions was measured. Signal density plots were generated using the BEDTOOLS(*8*) software, with data reported as reads per million reads mapped (RPM), and visualized using Integrative Genome Viewer (IGV) software (*10*).

### Assessment of inter-tissue correlations

At each CoRSIV, inter-tissue correlations of average proportional DNA methylation were computed for all tissue-pairs in which coverage requirements were satisfied in at least 10 individuals in both tissues. Pearson correlation was computed using the Python Scientific Library, with significance achieved at p<0.05. Inter-tissue correlation plots were visualized using the Python seaborn visualization library.

### CoRSIV/tissue DNA methylation clustering analysis

To assess the similarity of DNA methylation profiles across donors and tissues, donors with CoRSIV capture data in at least 4 tissues were considered. Next, CoRSIVs with sufficient coverage across all donors and tissues were selected. Finally, CoRSIV-average proportional DNA methylation values for each sample were clustered using the Euclidean distance metric and the average linkage method, and visualized using the seaborn Python visualization library.

### Cross-tissue analysis of gene expression vs. CoRSIV methylation

For each GTEx donor included in our analysis, tissue-specific gene expression profiles were downloaded from the GTEx data portal, expressed in transcripts per kilobase million (TPM). For each tissue, the analysis focused on CoRSIV associated genes expressed in that tissue (average TPM expression > 0.5). (Tibial nerve was not included in this analysis due to the small number of samples with RNA-seq data available.) Thyroid, lung, and cerebellum were considered ‘target tissues’. For each CoRSIV-associated gene expressed in each target tissue, we calculated the Pearson correlation between CoRSIV average methylation in that tissue and gene expression in that tissue. We then asked if the same correlation was found between CoRSIV average methylation in a ‘surrogate tissue’ (blood or skin) and gene expression in the target tissue.

Within each ‘expression tissue’ and ‘methylation tissue’ pair, p values were corrected for multiple hypothesis testing using the Benjamini Hochberg method, with significance achieved for adjusted p-value<0.05. Agreement of correlation between gene expression and DNA methylation between target tissue and surrogate tissues (statistically significant and in the same direction) was plotted in pie charts using GraphPad Prism. For specific CoRSIVs, scatterplots of tissue-specific gene expression vs. tissue-specific DNA methylation were generated using the seaborn Python visualization library.

### mQTL Analysis using CoRSIV capture data on GTEx Samples

Analysis of associations between CoRSIV-average DNA methylation and genetic variation in cis was performed using a previously described strategy relying on the Simes correction (*11*). Rather than test for all significant mQTL associations, this approach conservatively tests whether, at each CoRSIV, there is evidence of mQTL. For each donor, single nucleotide variant (SNV) profiles computed by the GTEx consortium were downloaded in vcf format (dbGaP accession phs000424.v8.p2). SNVs reported in dbSNP and with a minor allele frequency (MAF) of at least 5% were selected for further analysis. mQTL analysis was conducted independently for each tissue. For each CoRSIV, the number of donors with both sufficient coverage in the capture experiment for a specific tissue and with a WGS SNV profile available was determined; for each tissue, CoRSIVs with data for at least 20 donors were selected for mQTL analysis. To harmonize our mQTL analysis with those based on the Illumina BeadArray data, CoRSIV-average proportional DNA methylation values were converted to M-values (*12*) prior to analysis. Spearman rank correlation was computed for all SNVs within 1mb of each CoRSIV, using the EMatrixQTL R package (*13*), and the Simes correction was applied. Simes adjusted p-values for each CoRSIV were collected, and the false discovery rate (FDR) correction was applied across all CoRSIVs analyzed in each tissue, with significance achieved at FDR-adjusted p-value<0.05. The R^2^ variance explained by the linear model for each CoRSIV (in each tissue) was computed using the Python scientific library. For each significant mQTL association, a parametric analysis was carried out using using EMatrixQTL to determine the beta coefficient of the linear association between CoRSIV-average DNA methylation and the *cis* genetic variant.

Manhattan plots of mQTL associations were generated for each tissue and each CoRSIV using the R statistical system displaying all the mQTL candidates at p<0.001. Three-dimensional Manhattan plots of the significant mQTL associations across all CoRSIVS, capturing the distance between strongest associated SNV and CoRSIVs and the linear beta coefficient, were generated using the plotly R library. A distribution of the beta linear coefficients across all significant mQTL associations in each respective tissue was generated using the R library.

### Haplotype-based analysis using capture data on GTEx samples

SNVs reported in dbSNP and with a MAF of at least 5% were include in the haplotype-based analysis. PLINK 1.9 (*14, 15*) was used to identify haplotype blocks, with default parameters. Index SNVs were obtained by parsing the PLINK output for each individual block. Only CoRSIVs overlapping with haplotype blocks were considered for haplotype-based analysis. Further, only GTEx donors with a WGS profile were included, and within each tissue we considered only CoRSIVs with sufficient capture data on at least 20 donors. At each CoRSIV, the minor allele sum was computed across all the index SNVs for each donor, using the convention 0 for homozygous major allele, 1 for heterozygous SNV, and 2 for homozygous minor allele. Again, CoRSIV-average methylation values were transformed to M values. The Pearson correlation coefficient was computed between CoRSIV average DNA methylation and the minor allele sum of its overlapping haplotype block. Correction for multiple hypothesis testing was performed using the Benjamini-Hochberg correction, with significance achieved at FDR-adjusted p-value<0.05. Plots of DNA methylation at individual CoRSIVs vs. minor allele sum within overlapping haplotype blocks were generated using the Python scientific library.

### Analyzing consistency of CoRSIV mQTL across tissues

Recurrence of significant mQTL for each CoRSIV across the 6 tissues was assessed in two ways. First, at the most stringent level (the SNV level), an mQTL SNV-CoRSIV pair was considered recurrent if the same Simes-adjusted SNV was identified, and the beta coefficient had the same sign, within two or more tissues. Considering the high linkage disequilibrium among multiple SNVs within a haplotype block, we also evaluated recurrence at the haplotype block level. At this level, an mQTL association for a CoRSIV was considered consistent across multiple tissues if the Simes-adjusted SNVs identified in two or more tissues fell within the same haplotype block, and the beta coefficient had the same sign. mQTL recurrence was plotted as heat maps using the R statistical system.

### USC pediatric cohort – genotyping and CoRSIV capture bisulfite sequencing

Pediatric glioblastoma cases and controls were selected from the California Biobank, using information from the California Cancer and Vital Statistics registries. Cases were self-reported non-Latino whites born between 1982 and 2009, and subsequently diagnosed with glioblastoma (ICDO-3 code 9440). Controls were born in the same year with same gender and ethnic group as cases from anywhere in the state. Neonatal dried blood spots (approx 1.3 cm diameter) for each child were used for DNA extraction. DNA extraction, preprocessing and genotyping were performed as previously described (*16*). In brief, DNA was extracted from 1/3 of a dried blood spot with Genfind v3.0 (Beckman) reagents on an Eppendorf robot, followed by in house quality control procedures including nanodrop for purity and pico-green measurement for DNA quantity. Four hundred ng DNA was genotyped using the Affymetrix Axiom Precision Medicine Diversity Array (PMDA) at Thermo Affymetrix (San Jose CA), and SNP calls were extracted using Affymetrix Powertools. CoRSIV-capture bisulfite sequencing was performed using CoRSIV Capture v2.0 (Design ID: S3223244).

### USC Pediatric cohort CoRSIV capture data processing

For the USC pediatric whole blood cohort, Trim galore software was used for the quality control of the reads, which were aligned to hg38 genome using Bismark aligner (*9*). De-duplication was not carried according the Bismark guidelines for target capture sequencing. Bismark methylation extractor was used to do the methylation calling. CpG Methylation levels were averaged across CoRSIVs with at least 10x coverage.

### Independent analysis of USC pediatric samples for confirmation of CoRSIV mQTL and effects of local haplotype

CoRSIV capture bisulfite sequencing data on whole blood (newborn blood spots) were generated for 48 individuals from the USC pediatric cohort. One individual was removed from the analysis as a genetic outlier, leaving 47 samples for this analysis. Phased genotype data were generated, and SNVs reported in dbSNP and with a minor allele frequency (MAF) of at least 5% were selected for further analysis. CoRSIV-average DNA methylation values were converted to M-values (*12*) prior to analysis. Spearman rank correlation was computed for all SNVs within 1mb of each CoRSIV, using the EMatrixQTL R package (*13*), and the Simes correction was applied. Simes adjusted p-values for each CoRSIV were collected, and the false discovery rate (FDR) correction was applied across all CoRSIVs analyzed in each tissue, with significance achieved at FDR-adjusted p-value<0.05.

For the haplotype-based analysis, SNVs reported in dbSNP and with a MAF of at least 5% were included. PLINK 1.9 (*14, 15*) was used to identify haplotype blocks, with default parameters. Index SNVs were obtained by parsing the PLINK output for each individual block. At each CoRSIV, the minor allele sum was computed across all the index SNVs for each donor, using the convention 0 for homozygous major allele, 1 for heterozygous SNV, and 2 for homozygous minor allele. The Pearson correlation coefficient was computed between CoRSIV DNA methylation and the minor allele sum of its overlapping haplotype block. Correction for multiple hypothesis testing was performed using the Benjamini-Hochberg method, with significance achieved at FDR-adjusted p-value<0.05.

### Comparison of CoRSIV mQTL with HM450K mQTL results from GoDMC

GoDMC (Min et al. (*17*)) computed mQTL using 33,000 individuals with DNA methylation data generated in the HM450 platform. As described above, mQTL was calculated for the GTEx CoRSIV capture data using matrixEQTL software, regressing DNA methylation M-value against the genotype (0,1,2). To compare the summed total of mQTL detected at CoRSIVs vs. that reported by GoDMC, mQTL associations were identified with P < 10^-10^. This conservative P value was selected to avoid false positives, given the relatively small number of individuals in the GTEx CoRSIV analysis. With this cut-off, approximately 150,000 cis mQTL effects were detected in each study. Since methylation data were not available from GoDMC, methylation ranges (delta) were inferred by multiplying the slope of the linear model by 2 (x-axis of the genotype call). For each individual mQTL association, the product (delta)x(R^2^) yields the absolute quantity of interindividual methylation variation explained by the SNV. This metric was summed across all mQTL effects in each data set.

### Analysis of regional enrichments of transposable elements

To compare genomic enrichment of transposable elements flanking CoRSIVs vs. non-CoRSIV regions, repeat definitions encoded in the RepeatMasker track were downloaded from the UCSC genome browser build hg38. Repeats were analyzed at the level of repeat class, repeat class and repeat family and, finally, repeat class, repeat family, and repeat names. Only repeat sets with at least 10,000 entries were included in the enrichment analyses. CpG islands, defined by the UCSC genome browser on the human genome build UCSC hg38, were also downloaded. Analysis extended to +/-50,000 bp relative to each CoRSIV or comparison region. To compare the differential enrichment of one repeat subset R between two sets of genomic intervals A and B, we used BEDTOOLS (*8*) to determine repeat overlap between R and A or between R and B, within each of 50 genomic windows, cumulatively stepping by 1,000 bp increments from 0bp to 50,000bp. Within each window, an odds-ratio and p-value were computed using the Fisher’s exact test. Multiple testing correction was performed within each genomic window to adjust for the multiple tests performed at each repeat type, with significance achieved at an FDR-adjusted P < 0.01. Odds ratios (enrichments or depletions) surviving multiple testing correction were plotted across all repeat subsets and genomic windows using GraphPad Prism.

### Evolutionary selection analysis

Selection scores computed using Tajima’s D score (*18*) across CEU specimens profiled by the 1000 genomes project were downloaded (*19*). Selection scores compiled within a 30kb radius around CoRSIVs, control regions, tDMR regions, or 450k probes were plotted using the R statistical analysis system.

### Enrichment of GWAS trait SNVs

We employed permutation testing to determine the extent to which CoRSIV mQTL SNVs (Simes SNVs) are enriched for trait-associated SNVs from the NHGRI GWAS catalogue (downloaded October 2020). For each of 8 manually curated trait categories (*20*), we generated a null distribution by randomly selecting one SNV from the GTEx database (MAF >= 0.05) within 1 Mb up- or downstream from the center of each CoRSIV. We then determined whether this randomly chosen SNV overlapped an NHGRI trait-associated SNV from that category. This process was repeated 10,000 times to yield a null distribution for each trait category. The numbers of actual overlaps between CoRSIV mQTL SNVs and NHGRI trait-associated SNVs were compared to these null distributions using one-proportion Z tests. Bonferroni-adjusted p-values for these tests are reported. Enrichment was defined as the number of actual overlaps between CoRSIV mQTL SNVs and NHGRI trait-associated SNVs divided by the mean of the null distribution.

### CoRSIV +/- 20kb SNVs heritability enrichment with/without controlling for 53 baseline features

To evaluate heritability for a variety of traits (i.e., cancer and metabolic diseases) in CoRSIV +/-20kb SNVs, we used stratified linkage disequilibrium score regression (s-LDSC) (*21*) . First, the targeted CoRSIV regions were lifted to hg19 using the UCSC “liftover” software, and “bedtools” software was used to add the +/- 20kb flanking. CoRSIV +/- ‘20kb’ distance was used because the majority of significant mQTL occurred within these regions. European population plink files used by LDSC software (v. 1.0.1) were downloaded from the 1000 Genomes project.

In step 1, “make_annot.py” python script available in LDSC software was used to generate SNV annotation for CoRSIV +/- 20kb and 53 ‘baseline’ (*21*) bed files, creating (.annot) files typically consisting of CHR, BP, SNP, and CM columns, followed by one column per annotation, with the value of the annotation for each SNP (0/1 for binary categories). Two separate sets of annotation files were generated for CoRSIV +/- 20kb with and without 53 ‘baseline’ features.

In step 2, two annotation file sets generated in step1 were used to estimate annotation-specific LD scores using 1000 Genomes phase3 plink files and 1000G HapMap3 SNPs excluding MHC region in chr6 (according to the LDSC user manual). LDSC.py script with the following parameters suggested by the manual was used. --bfile flag points to the plink format fileset; The --l2 flag tells ldsc to compute LD Scores. The --ld-wind-cm flag tells lsdc to use a 1 cM window to estimate LD Scores.

In step 3, to calculate partitioned heritability for different traits, summary GWAS statistics were downloaded for 4 cancer and 12 metabolic disease categories from (https://alkesgroup.broadinstitute.org/LDSCORE/all_sumstats/). The LDSC.py script with --h2 flag compute partitioned heritability and outputs enrichment scores, for CoRSIV +/- 20kb regions (with/without controlling for 53 baseline features) for each trait.

---

**Fig. S1.**
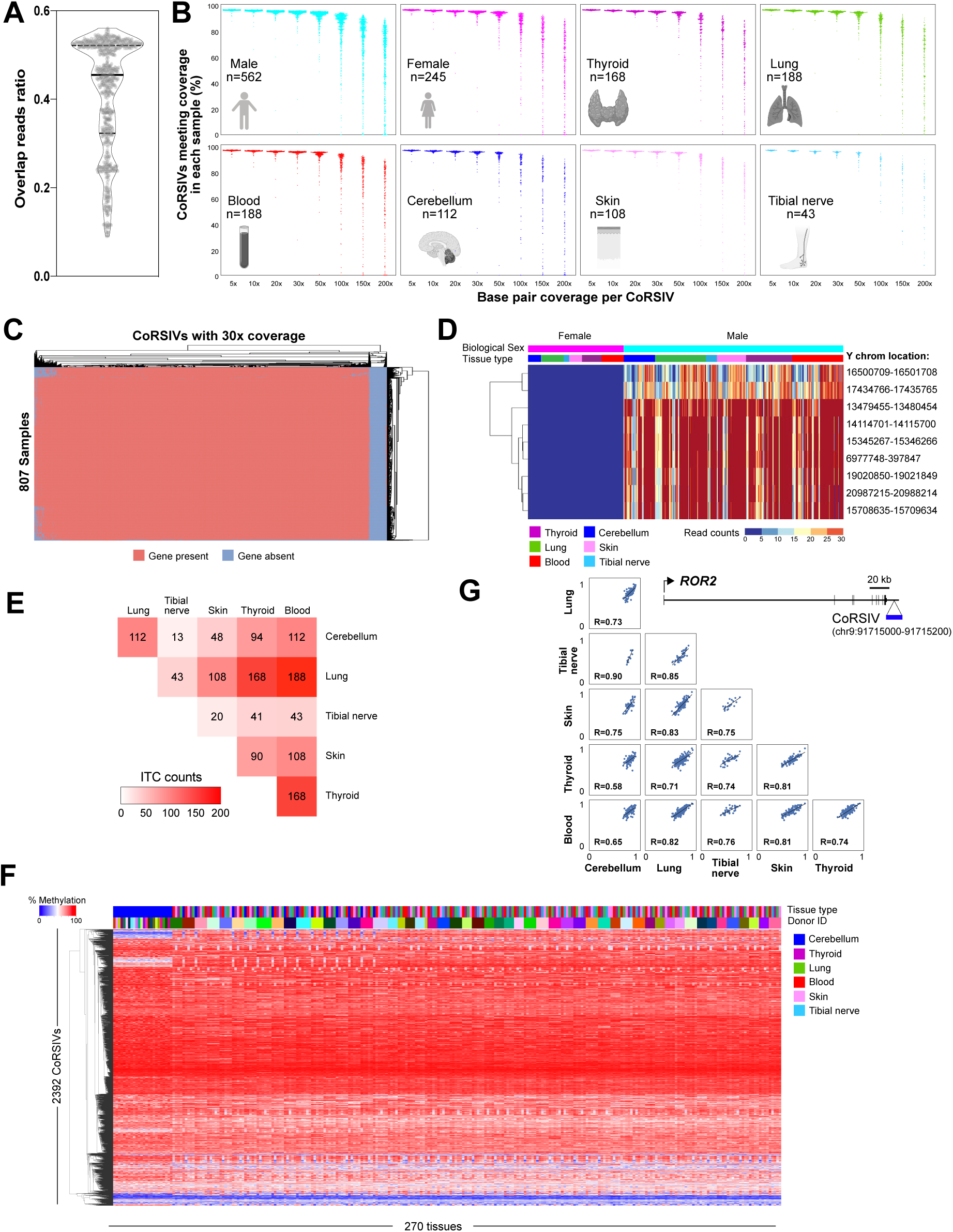
CoRSIV capture efficiency, quality control, and inter-tissue correlation in DNA methylation. **(A)** Violin plot shows, for each of 807 samples, the proportion of reads that were on target (i.e. completely within a target region). **(B)** Percentage of CoRSIVs for which target-capture bisulfite sequencing achieved various read depths, by sex, and by tissue type. Each point represents a sample; numbers of samples are shown. **(C)** Dichotomous heat map showing which of 4,483 CoRSIVs (columns) are covered at ≥30x depth in each of 807 samples (rows). A small fraction of CoRSIVs proved difficult to capture across all samples. **(D)** Read-depth across a panel of Y-chromosome probes confirms correct biological sex for all 807 samples (quality control). **(E)** Heat map shows numbers of samples available to calculate inter-tissue correlations. (**F**) For the 270 tissue samples from 53 donors with data on at least 5 tissues (including cerebellum), unsupervised hierarchical clustering of methylation data at 2,340 fully informative CoRSIVs organizes mainly by donor, but also forms a minor cerebellum cluster (left hand side). (**G**) Inter-tissue correlation plots for a CoRSIV near *ROR2* show that, despite higher methylation in cerebellum relative to other tissues, high inter-tissue correlations are maintained.

**Fig. S2.**
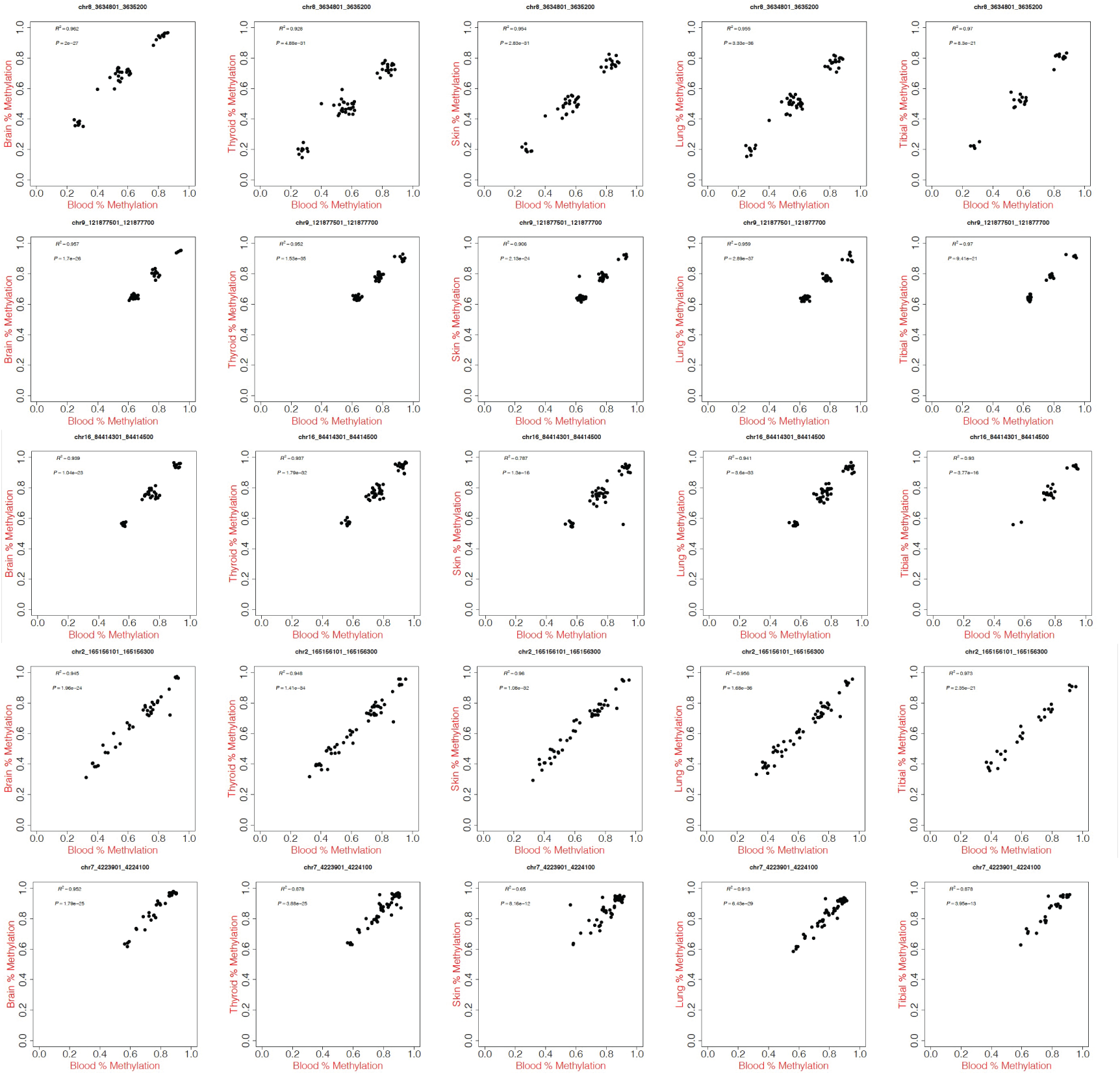
Inter-tissue correlation (ITC) plots show that methylation in blood is associated with that in brain, thyroid, skin, lung, and tibial nerve. Each row shows data on one CoRSIV. The first three and last two rows show CoRSIVs with three modes of methylation, and a uniform distribution, respectively (the most common patterns observed). For ITC plots on all the CoRSIVs see (five tissues vs. blood).

**Fig. S3.**
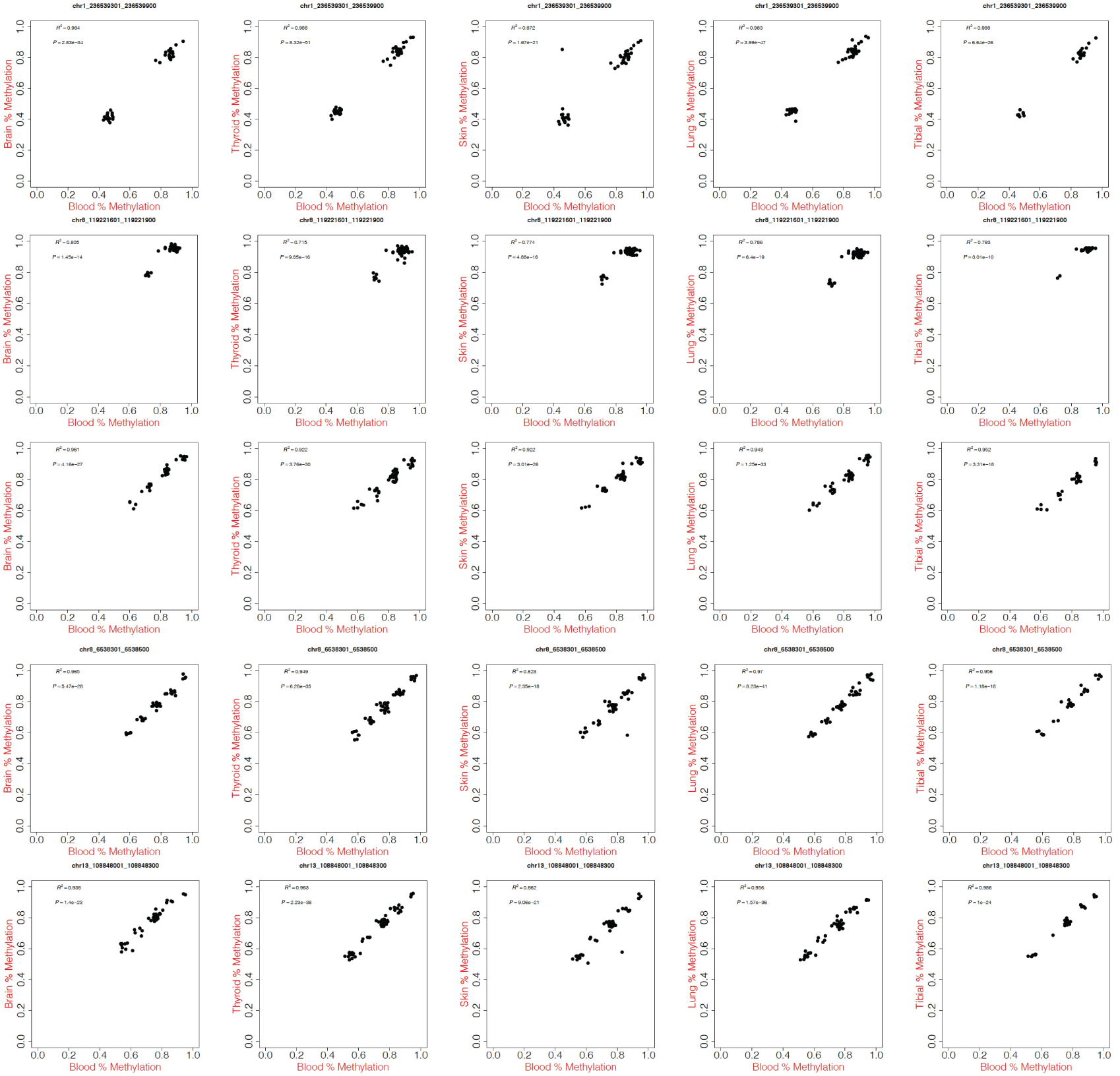
Inter-tissue correlation (ITC) plots show that methylation in blood is associated with that in brain, thyroid, skin, lung, and tibial nerve. Each row shows data on one CoRSIV. These plots show examples of infrequently observed patterns including two, four, or five discrete modes. For ITC plots on all the CoRSIVs see (five tissues vs. blood).

**Fig. S4.**
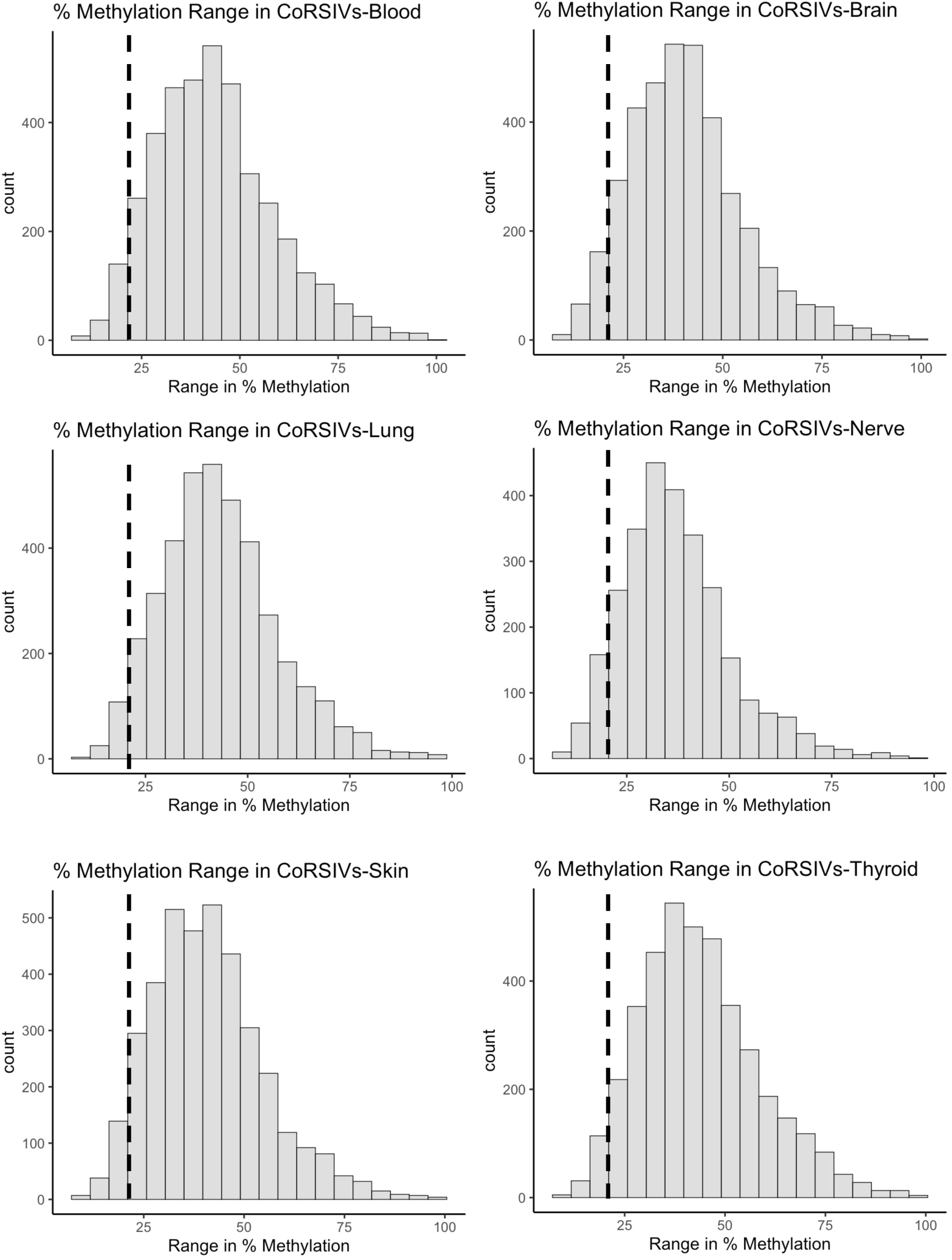
CoRSIVs exhibit high interindividual variation in every tissue examined. Each plot shows the distribution of interindividual methylation range across 4,086 CoRSIVs. In all six tissues, nearly all CoRSIVs show an interindividual range > 20% (dashed line).

**Fig. S5.**
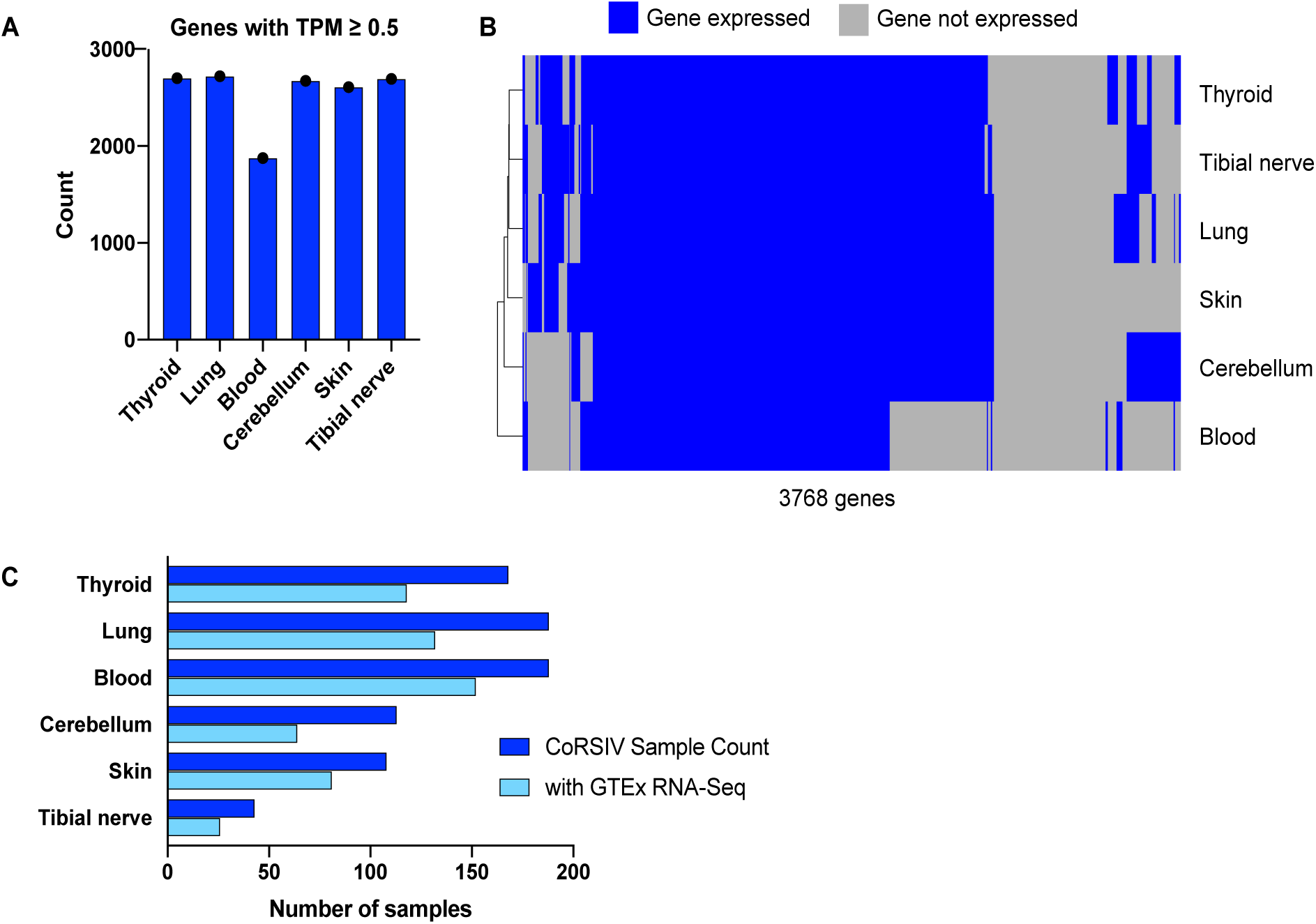
Supporting data for cross-tissue analysis of methylation and gene expression. **(A)** Numbers of CoRSIV-associated genes expressed (transcripts per million (TPM) >= 0.5) in each tissue type. **(B)** Heat map of expressed genes across the six tissues. **(C)** Except for tibial nerve, both methylation and gene expression data were available for more than 60 individuals.

**Fig. S6.**
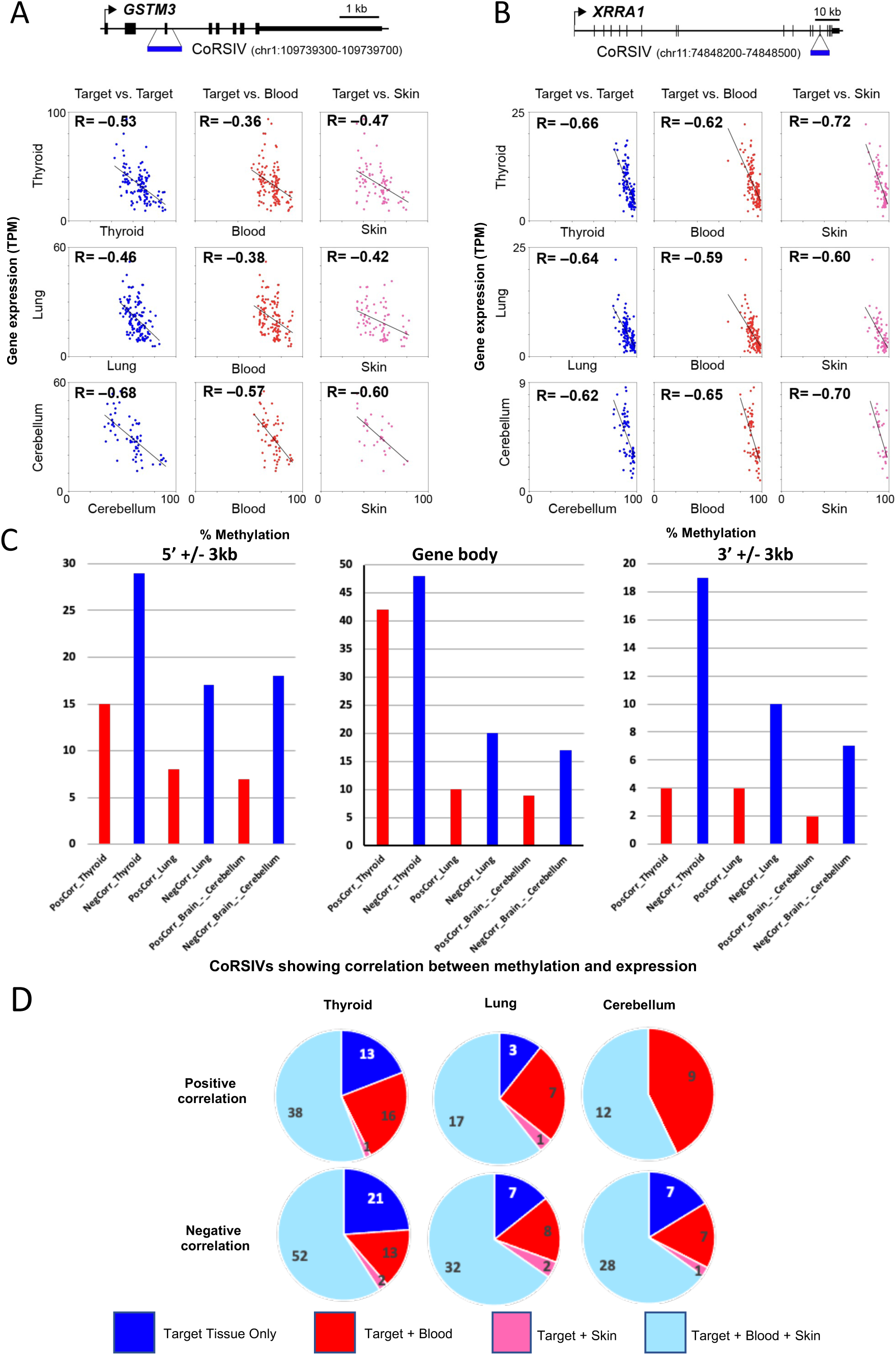
CoRSIV methylation in blood or skin predicts expression of associated genes in less accessible tissues. (**A)** Within- and between-tissue correlations of *GSTM3* expression vs. DNA methylation at a CoRSIV within *GSTM3*. (**B)** Within- and between-tissue correlations of *XRRA1* expression vs. DNA methylation at a CoRSIV within *XRRA1*. **(C)** Number of CoRSIVs with positive and negative correlations in three tissues according to location of CoRSIV with respect to transcription start site (5’), gene body, and transcription end site (3’). (**D)** Thyroid, lung, and cerebellum are considered target tissues, and blood and skin are surrogate tissues. Across all CoRSIV-gene pairs with significant correlations between CoRSIV methylation and expression of an associated gene in a target tissue, over 75% show the same correlation when methylation in blood is used as surrogate; using skin as surrogate, over 60% do.

**Fig. S7.**
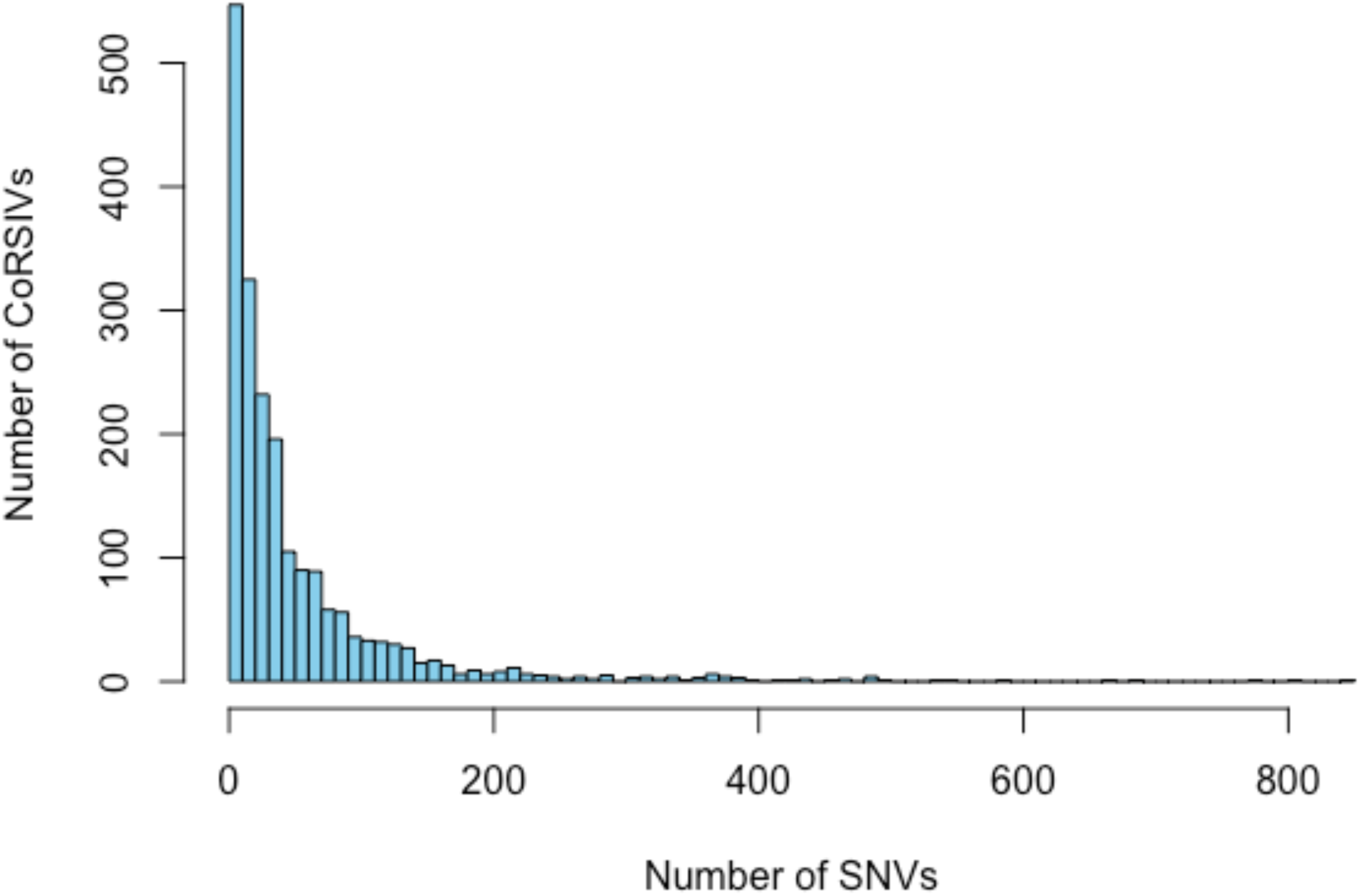
Histogram of the number of genetic variants (SNVs) influencing methylation at each CoRSIV (mQTL P <10^-10^).

**Fig. S8.**
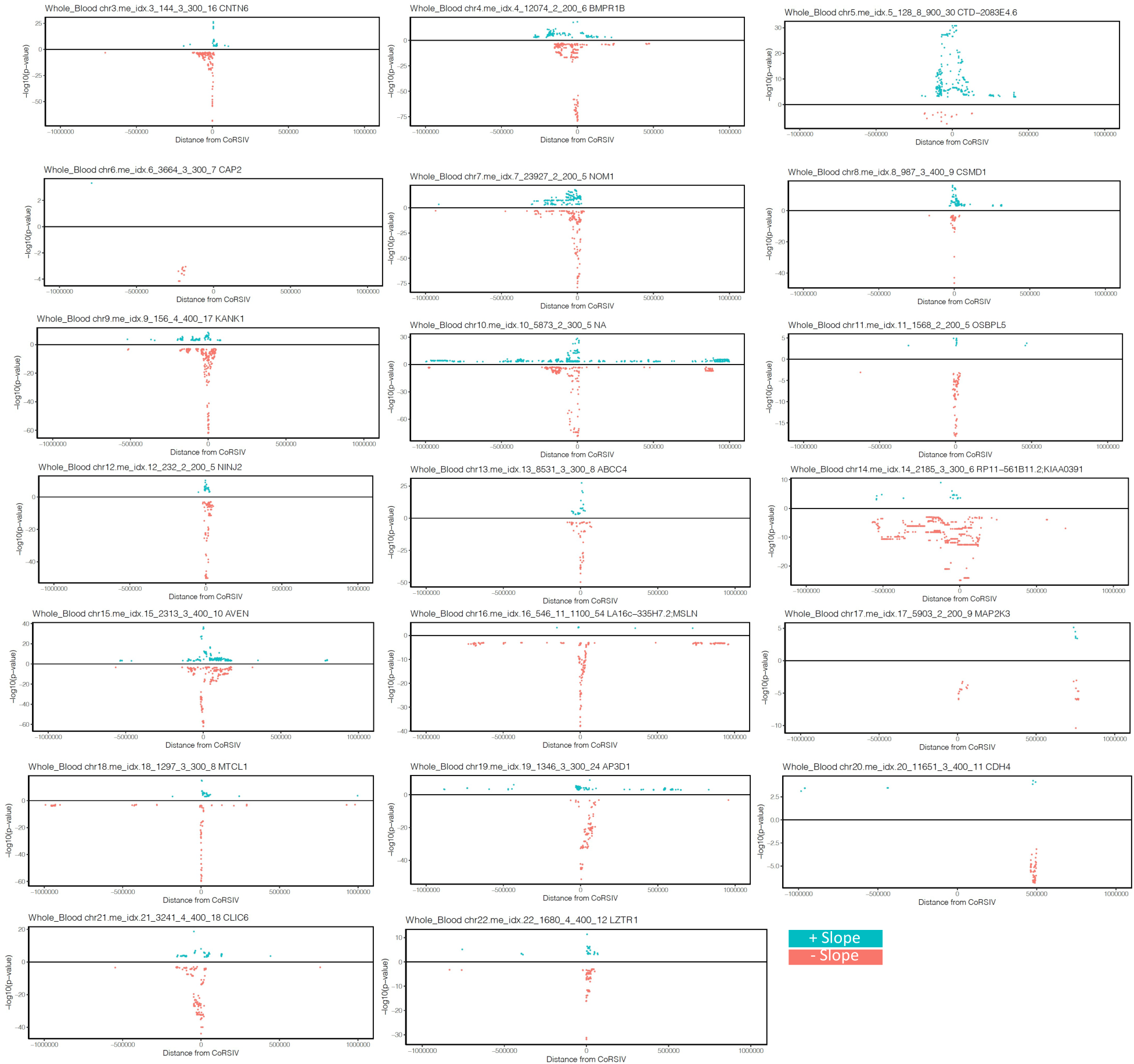
Representative Manhattan plots of associations at individual CoRSIVs, in blood. Significant associations are shown for all SNVs within 1Mb of each CoRSIV; positive and negative beta coefficients are plotted in blue and red, respectively.

**Fig. S9.**
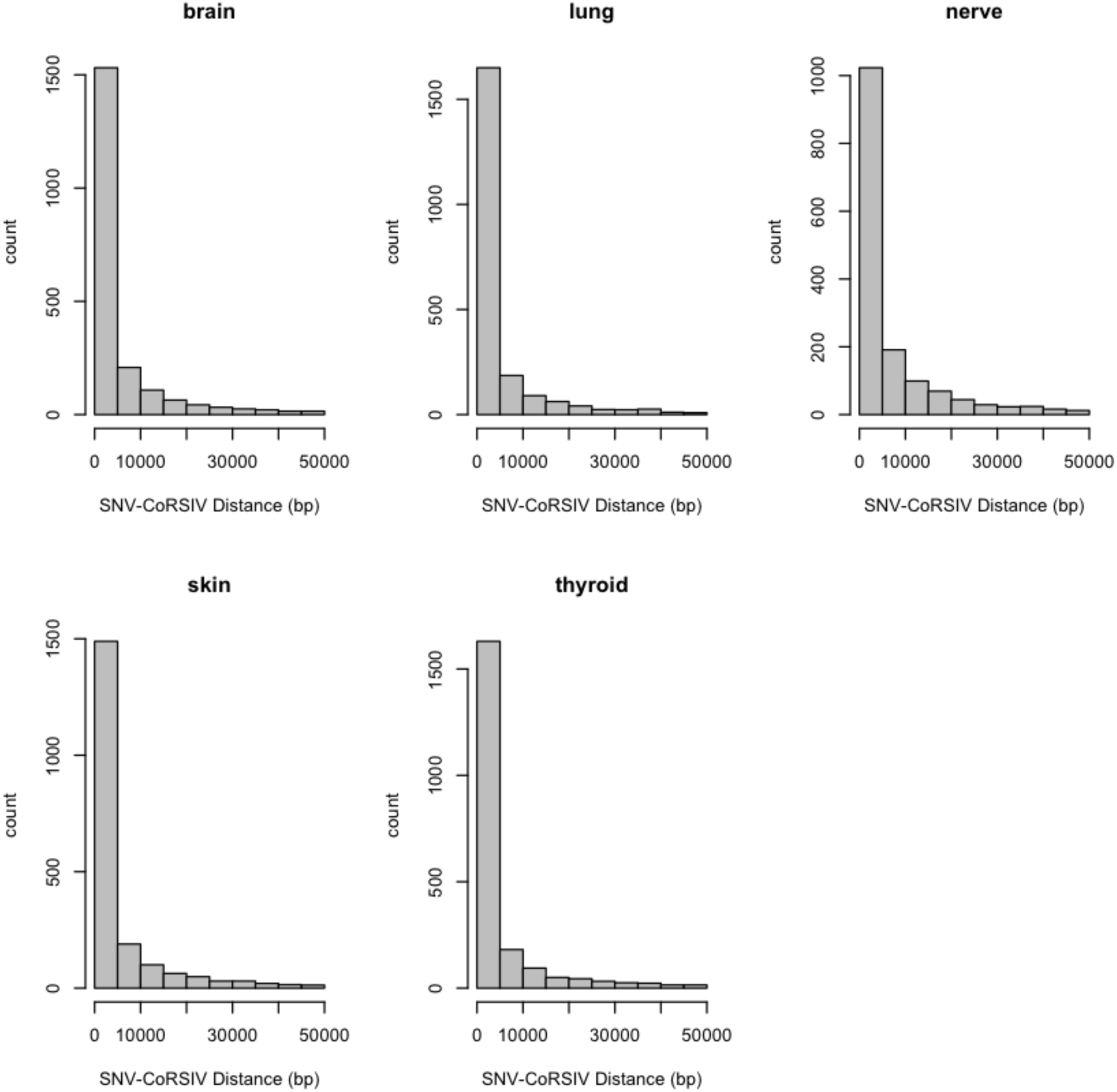
Distribution of distances between CoRSIVs and corresponding Simes SNVs, in five different tissues. All appear nearly identical to the distribution in blood (Fig. 2C). The majority of Simes SNVs are within 10 kb of each CoRSIV.

**Fig. S10.**
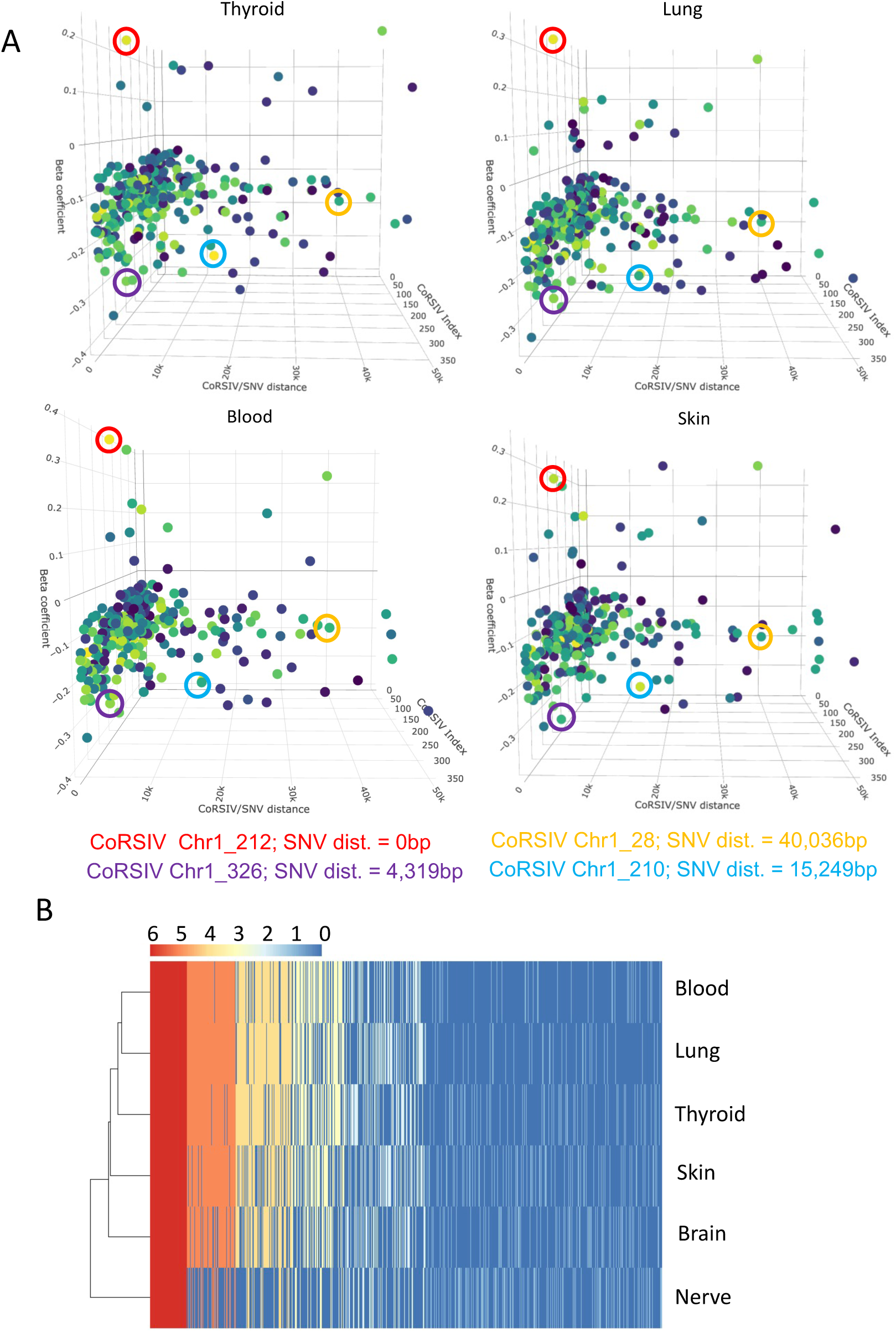
Concordance of mQTL effects across different tissues. **(A)** Three-dimensional Manhattan plots for chromosome 1 show four examples (circled) of CoRSIVs at which the exact same SNV was independently identified as the Simes SNV in thyroid, lung, blood, and skin. **(B)** For each of 4,086 CoRSIVs, heat map depicts the number of tissues in which the same SNV is identified as the Simes SNV.

**Fig. S11.**
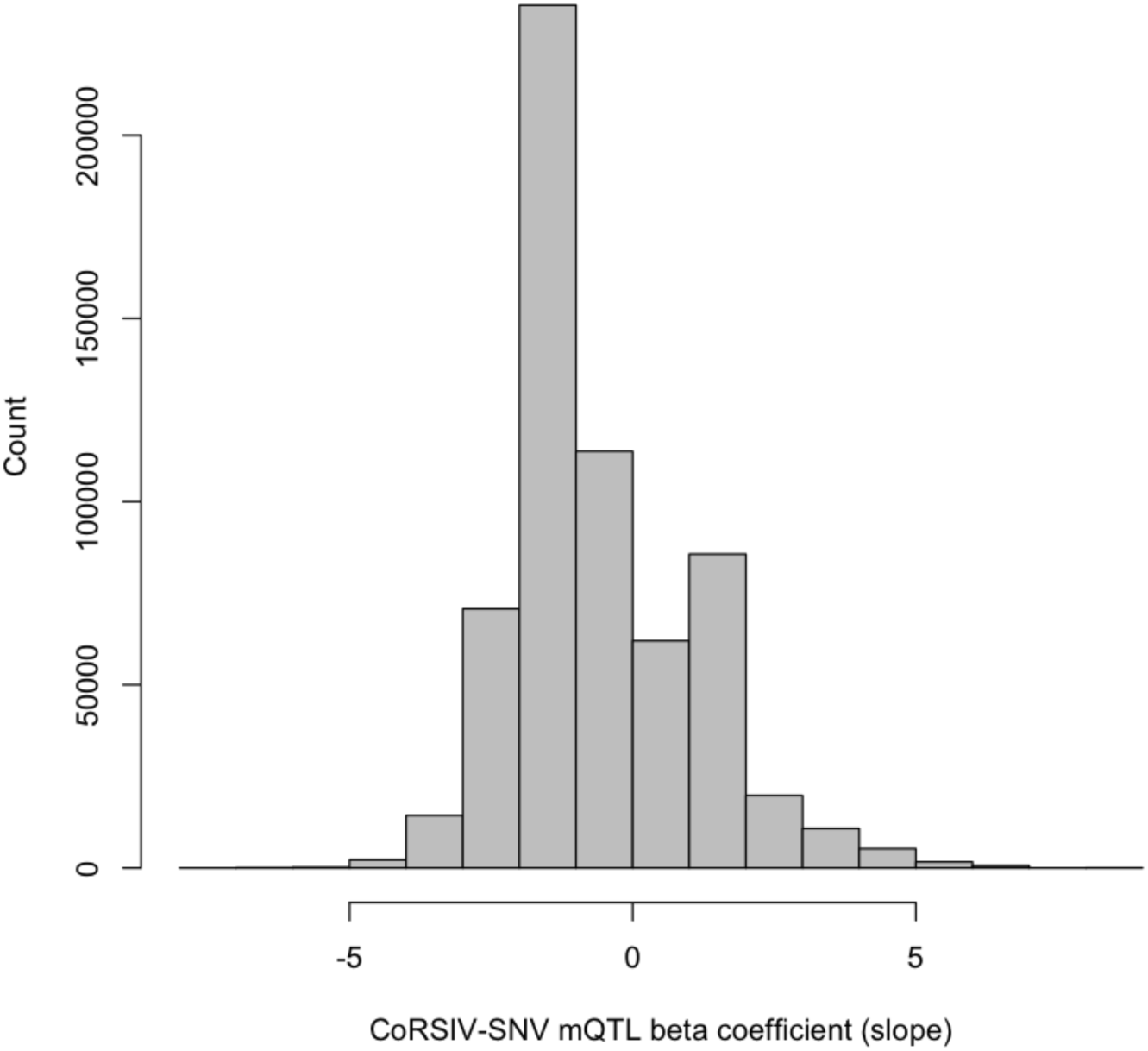
All mQTL associations at CoRSIVs are biased toward negative beta coefficients. Shown is the distribution of all 146,698 mQTL associations detected within 1Mb of CoRSIVs (P < 10^-10^) in blood. Similar to that of Simes SNVs (Fig. 2F), the bias toward negative beta coefficients is obvious.

**Fig. S12.**
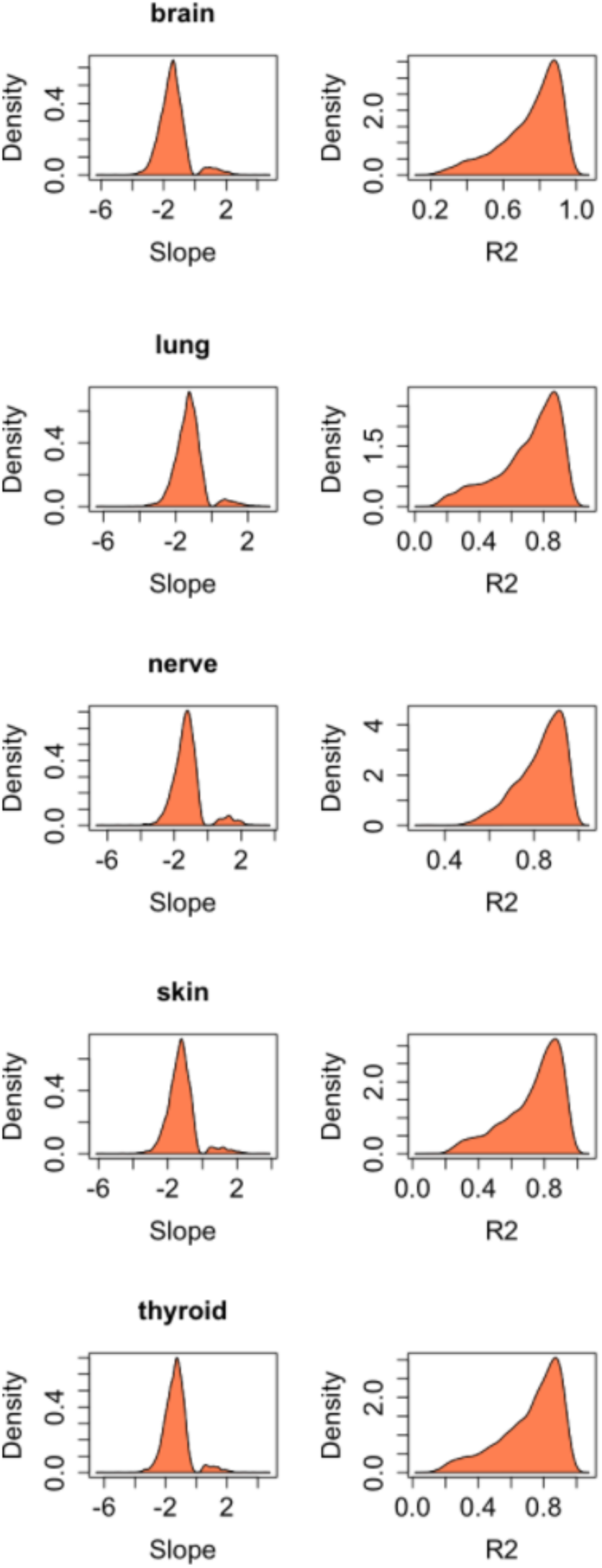
Distribution of Simes mQTL slope (i.e. beta coefficient) and R^2^ (goodness of fit) for brain, lung, tibial nerve, skin and thyroid are strikingly similar to those obtained for blood (Fig. 2 F, I).

**Fig. S13.**
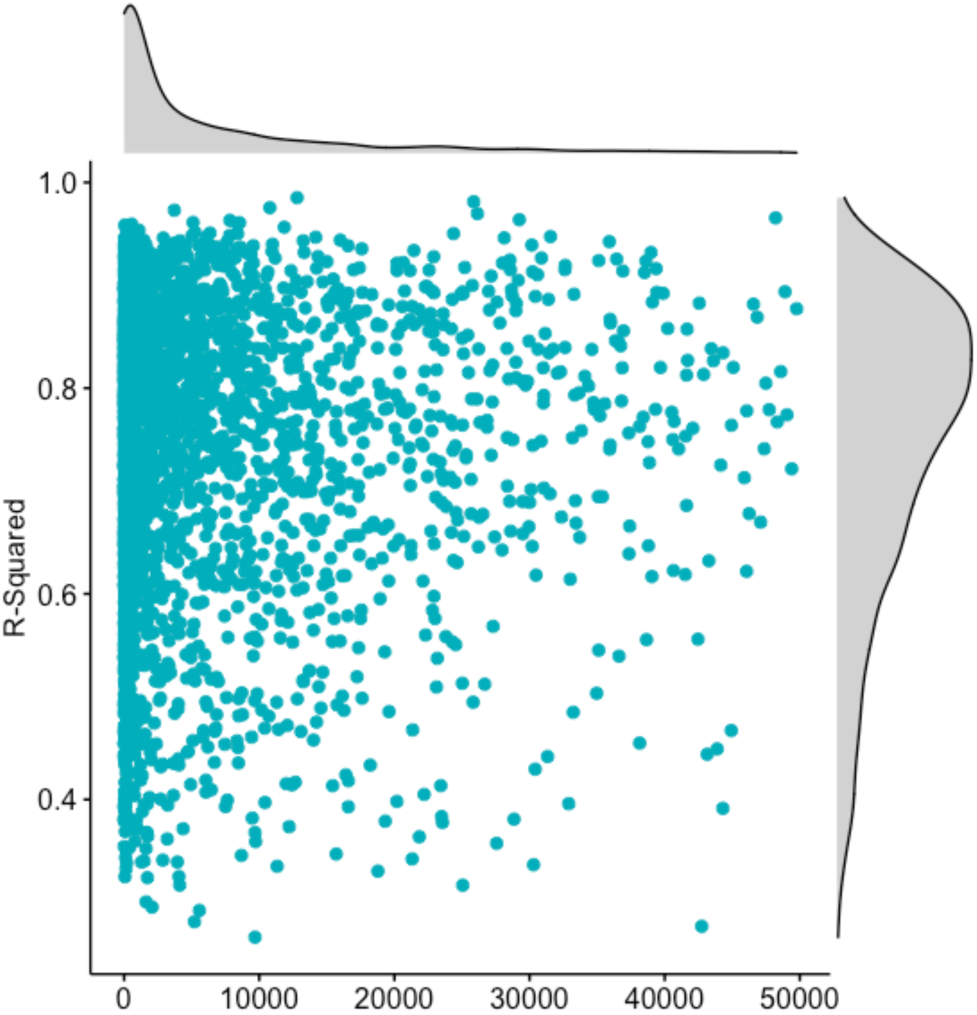
Scatter plot of R-squared (goodness of fit) vs. CoRSIV-SNV distance for Simes mQTLs. The bias toward high R^2^ mQTL effects is observed even at considerable CoRSIV-SNV distances.

**Fig. S14.**
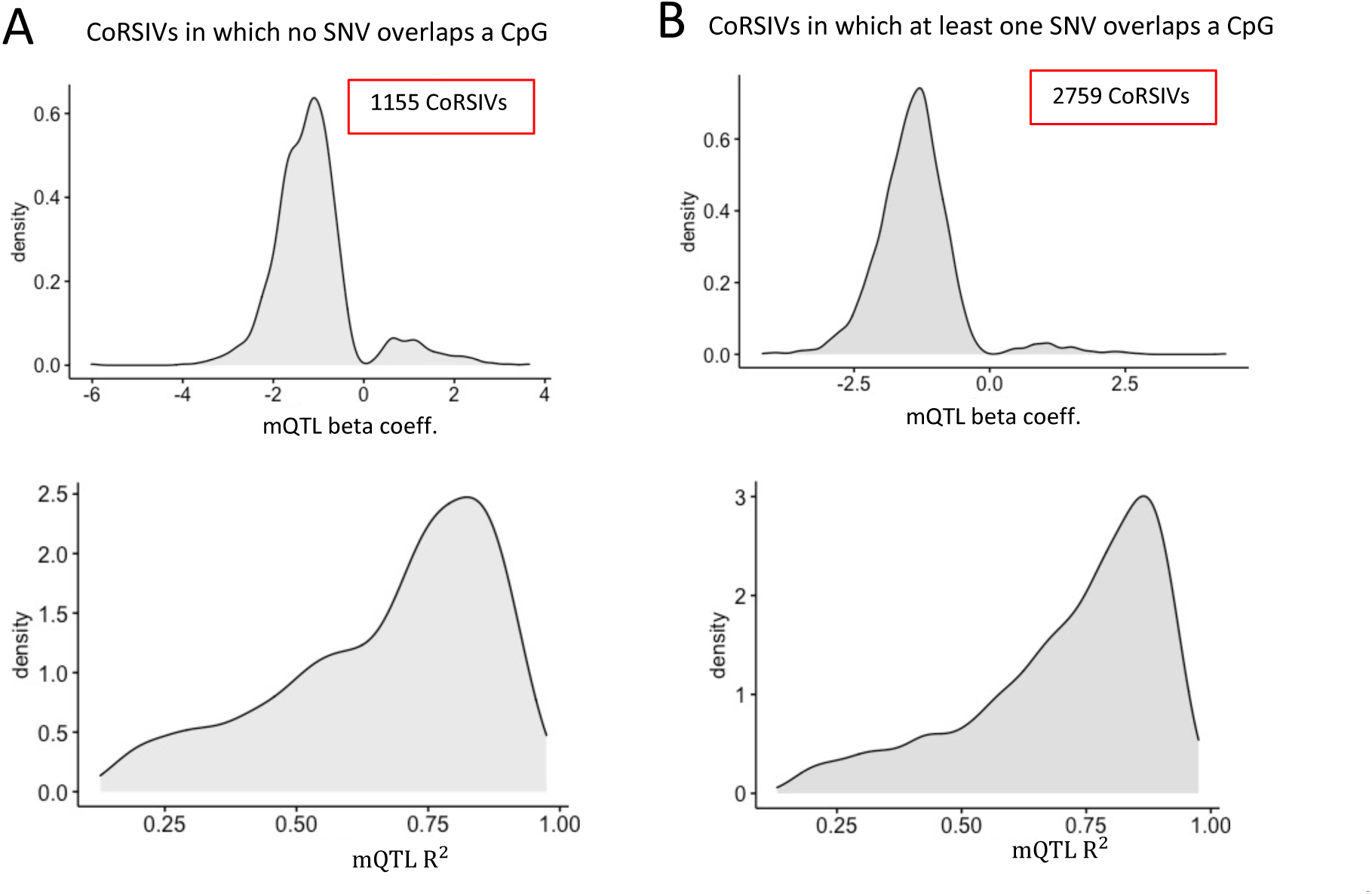
Neither the bias toward negative beta coefficients nor the tendency for high R^2^ of mQTL effects at CoRSIVs is explained by SNVs overlapping CpGs within CoRSIVs. (A) The set of 1155 CoRSIVs with no such overlaps. (B) The set of 2759 CoRSIVs for which at least on SNV overlaps a CpG within the CoRSIV. In both cases, the biases toward negative beta coefficients and high R^2^ are observed.

**Fig. S15.**
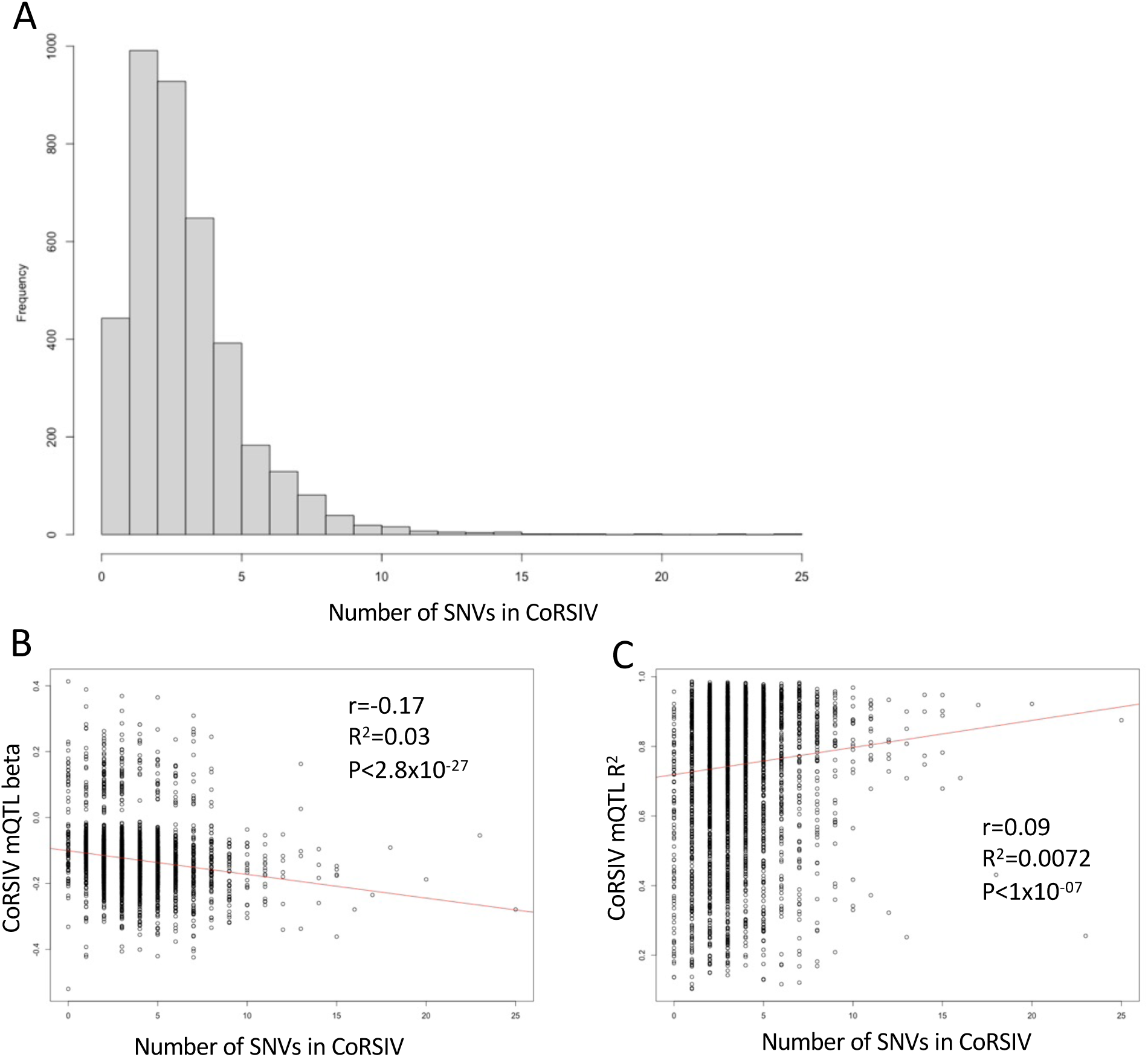
SNVs within CoRSIVs do not explain either the bias toward negative beta coefficients or the strong R^2^ of mQTL effects at CoRSIVs. (A) Distribution of the number of SNVs detected within each of 4,086 CoRSIVs. (B) CoRSIV mQTL beta coefficient is only weakly associated with the number of SNVs in each CoRSIV. (C) CoRSIV mQTL R^2^ is only weakly associated with the number of SNVs in each CoRSIV.

**Fig. S16.**
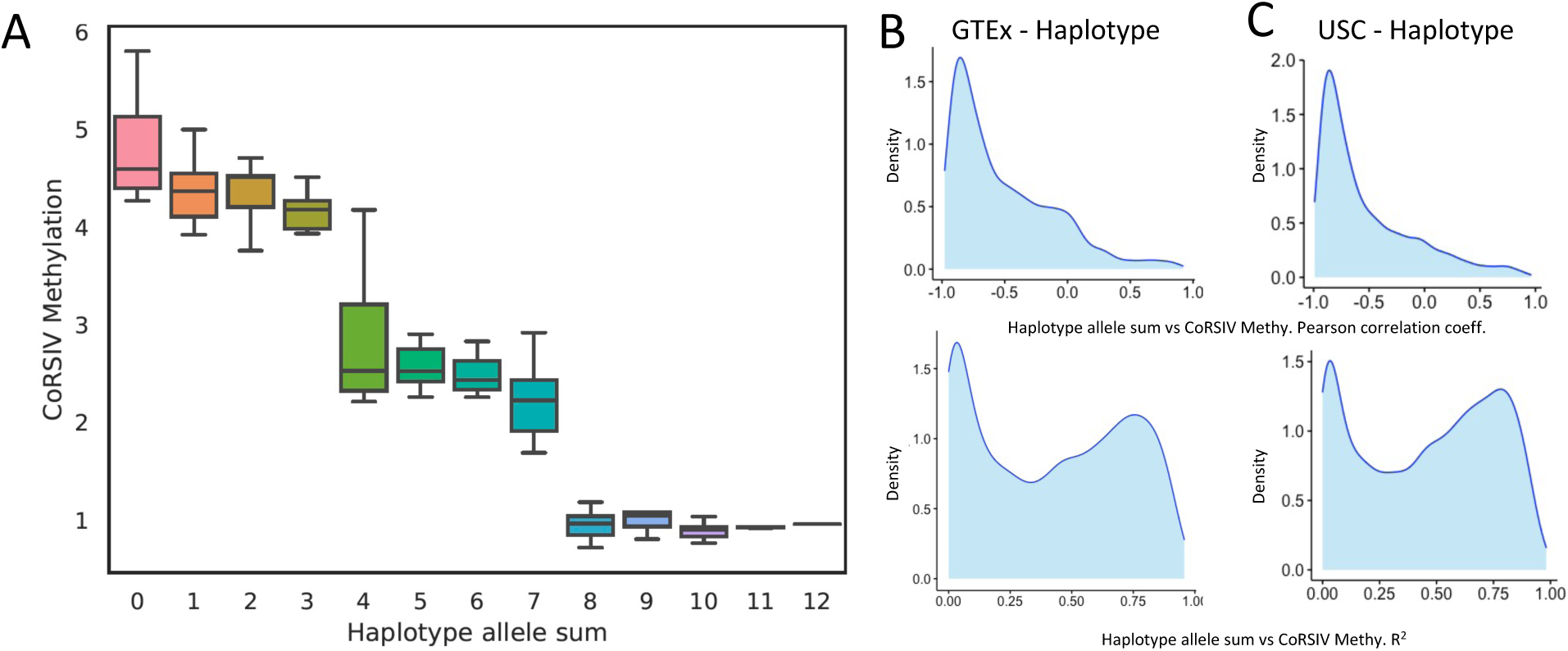
Haplotype-based approach to assess local genetic influences on CoRSIV methylation. **(A)** Example of association of average methylation in blood at one CoRSIV (chr22:50677397-50683133) vs. haplotype allele sum (sum of minor alleles in each individual) for its overlapping haplotype block. Shown are data on 170 individuals grouped by haplotype allele sum. **(B)** Top: Distribution of Pearson correlation coefficients for all such associations (CoRSIV methylation vs. haplotype allele sum) across 4,471 CoRSIVs assessed in 188 GTEx donors. As in the mQTL analysis (Fig. 2F), negative coefficients predominate. Bottom: Distribution of R^2^ (goodness of fit) across all such associations in GTEx donors. Local haplotype explains much of the variance in methylation. **(C)** Independent corroboration in USC cohort. Top: Distribution of Pearson correlation coefficients for all such associations (CoRSIV methylation vs. haplotype allele sum) across 4,471 CoRSIVs studied in 47 newborns in USC cohort. Bottom: Distribution of R^2^ across all such associations in USC cohort.

**Fig. S17.**
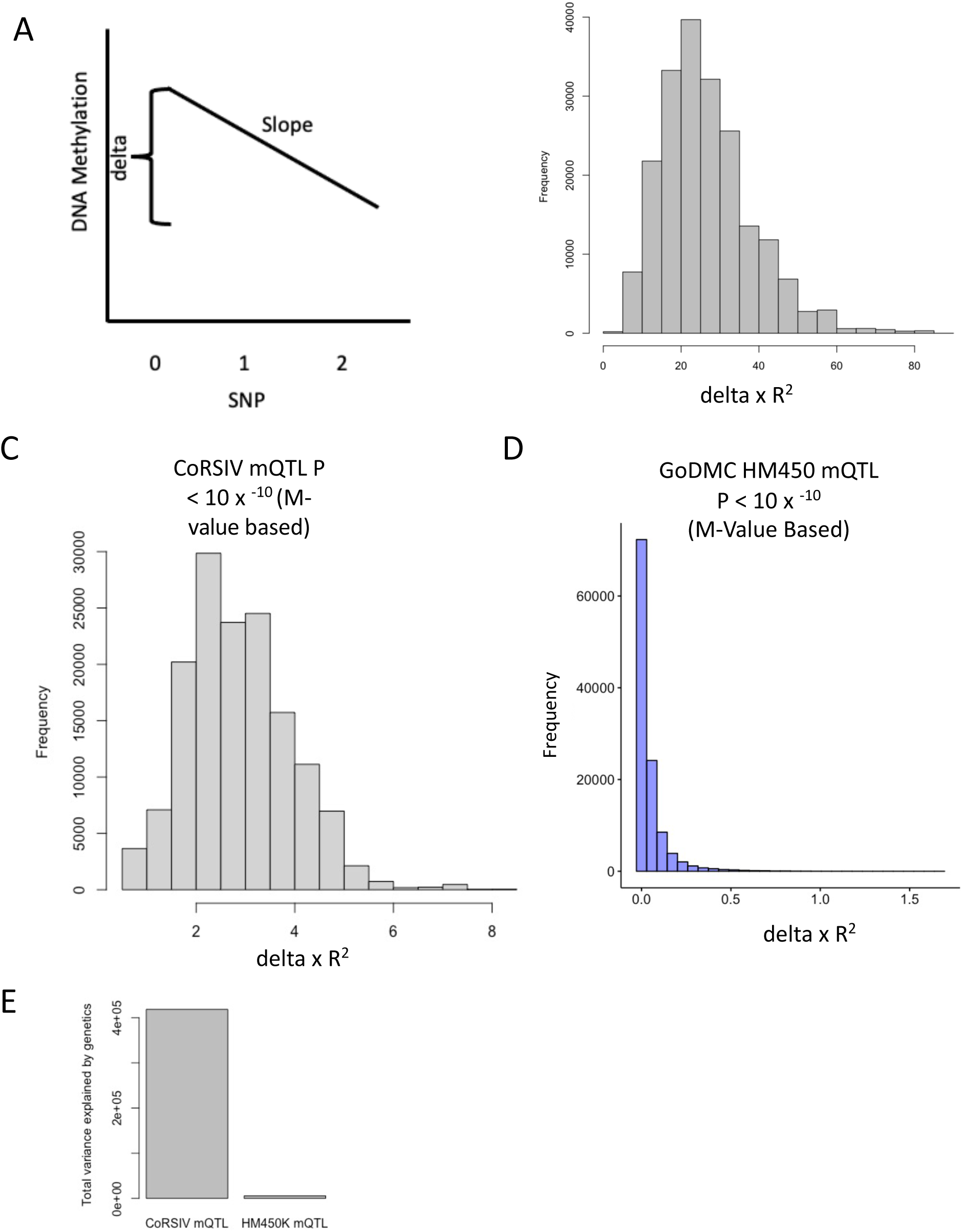
Approach for comparing total quantity of mQTL across different data sets. **(A)** With exactly 3 possible genotypes at every SNV, the methylation difference (delta) associated with each SNV can be easily determined from the beta coefficient (slope) of the mQTL association. **(B)** The product (delta)x(R^2^) measures the total amount of variation in methylation that is determined by the SNV genotype. Distribution of (delta)x(R^2^) for all 146,698 mQTL associations (P < 10^-10^) across 2,738 CoRSIVs in blood of 188 individuals. **(C)** The same distribution, recalculated using M-values. **(D)** Distribution of (delta)x(R^2^) for all 154,527 mQTL associations (P<10^-10^) detected using the HM450 platform to study blood of 33,000 individuals (GoDMC data – Min et al 2021 Nat. Genetics). Although the calculations were performed the same way as in (C), note the very different x-axis scales. **(E)** Area under the curve in (C) vs. that in (D). The summed total variance in methylation explained by cis mQTL at CoRSIVs is 72 fold greater than that detected in the GoDMC report.

**Fig. S18.**
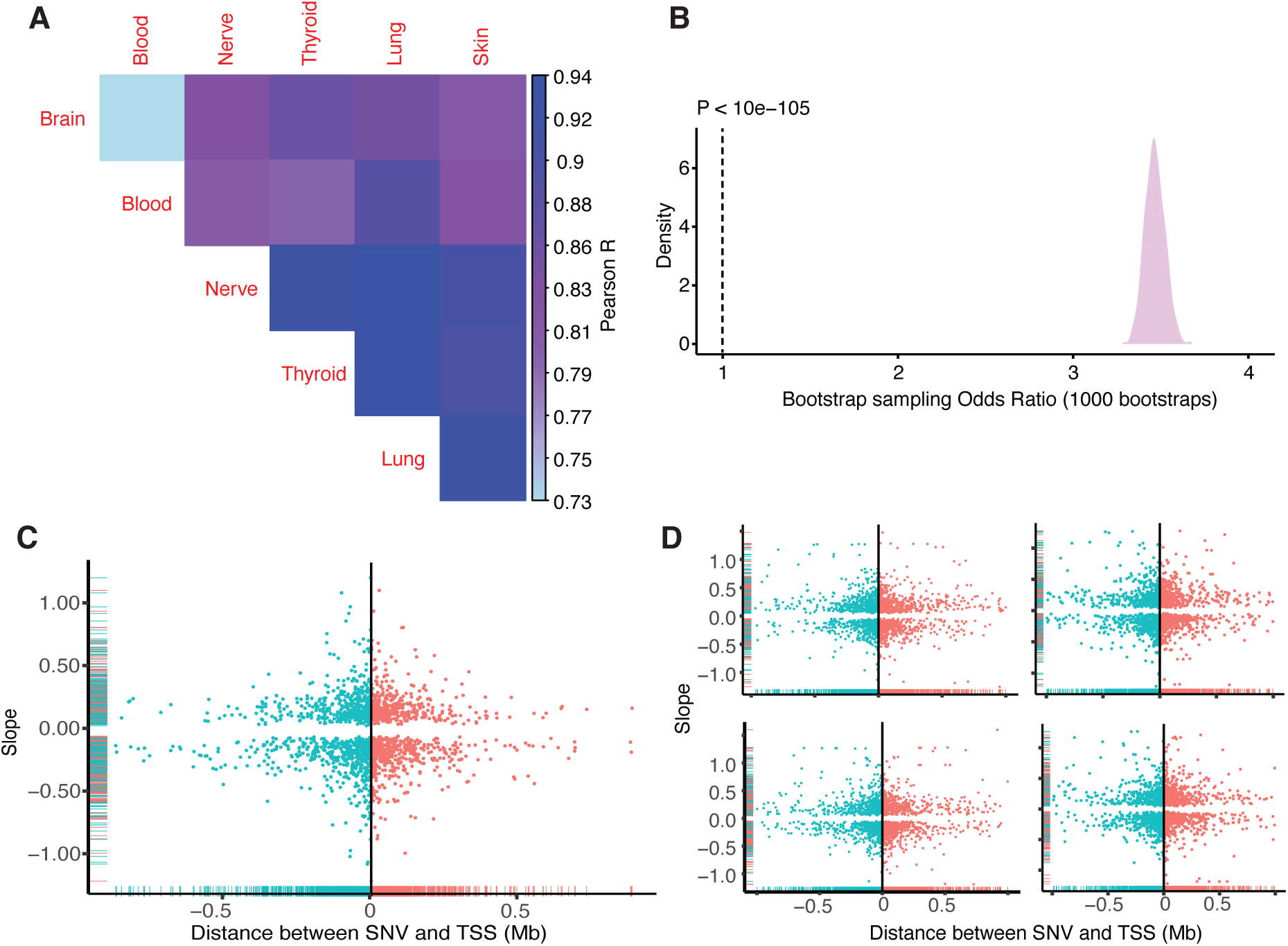
Characteristics of eQTLs overlapping Simes SNVs. **(A)** Pearson correlation between eQTL slopes across 5 tissues. **(B)** Enrichment of Simes SNVs in eQTL compared to bootstrapped eQTLs from CoRSIV +/- 50kb flanking regions. Fisher test P-values < 1×10^-105^. **(C)** eQTL slope vs. distance between SNV and TSS for Simes SNVs. **(D)** eQTL slope vs. distance between SNV and TSS for bootstrapped eQTLs (four representative plots from 1000 bootstraps are shown).

**Fig. S19.**
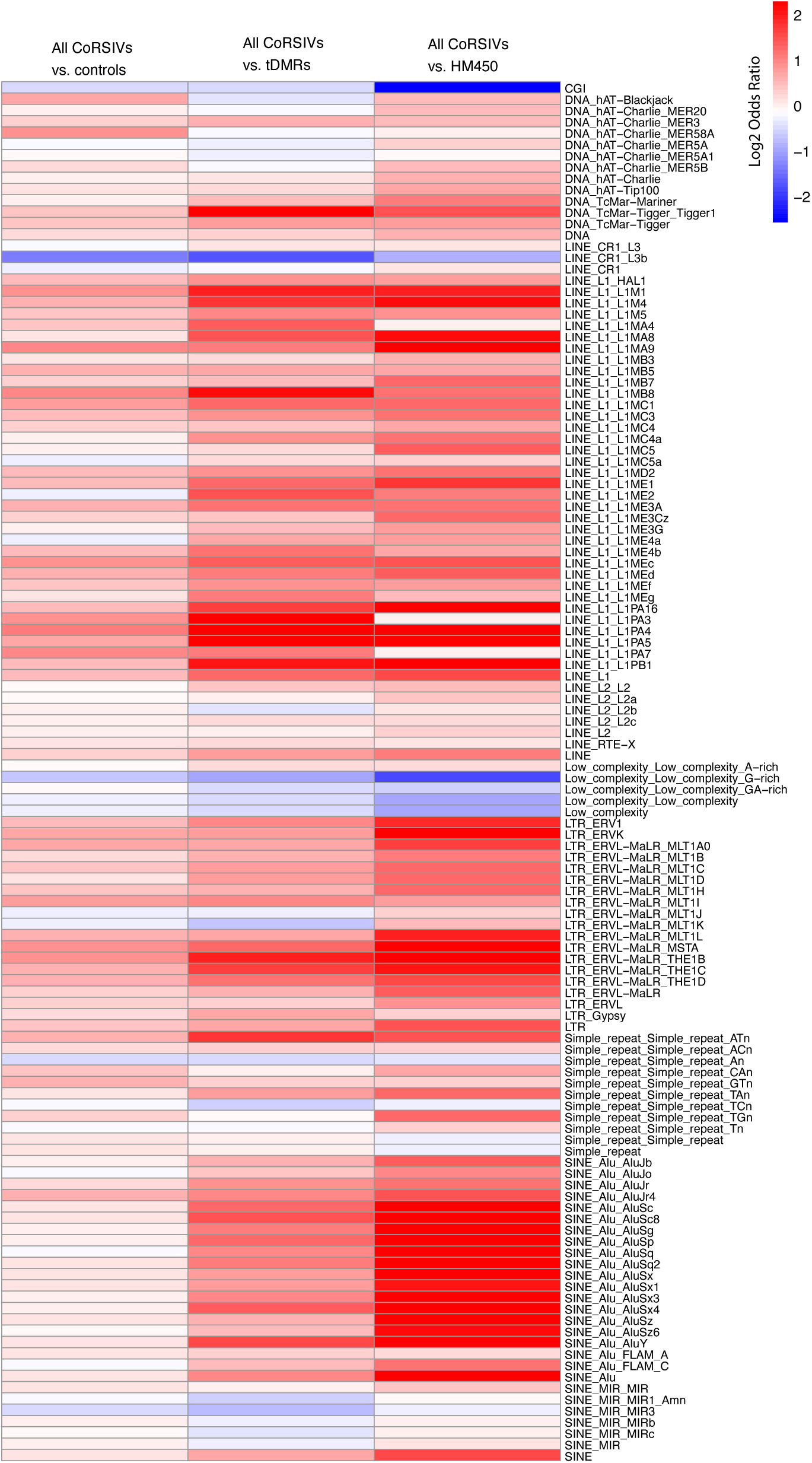
Overlap of transposable elements over all CoRSIV regions compared to controls, tDMRs and HM450.

**Fig. S20.**
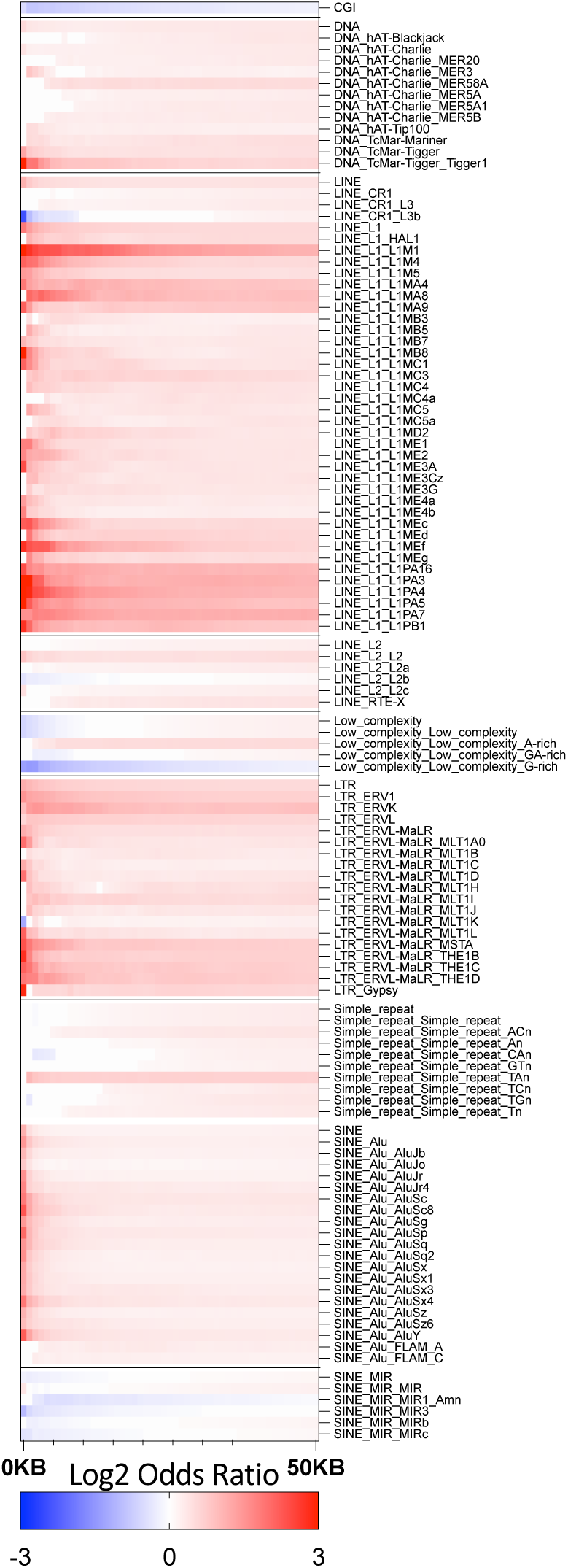
Transposable element enrichment for Genic CoRSIVs vs. tDMRs.

**Fig. S21.**
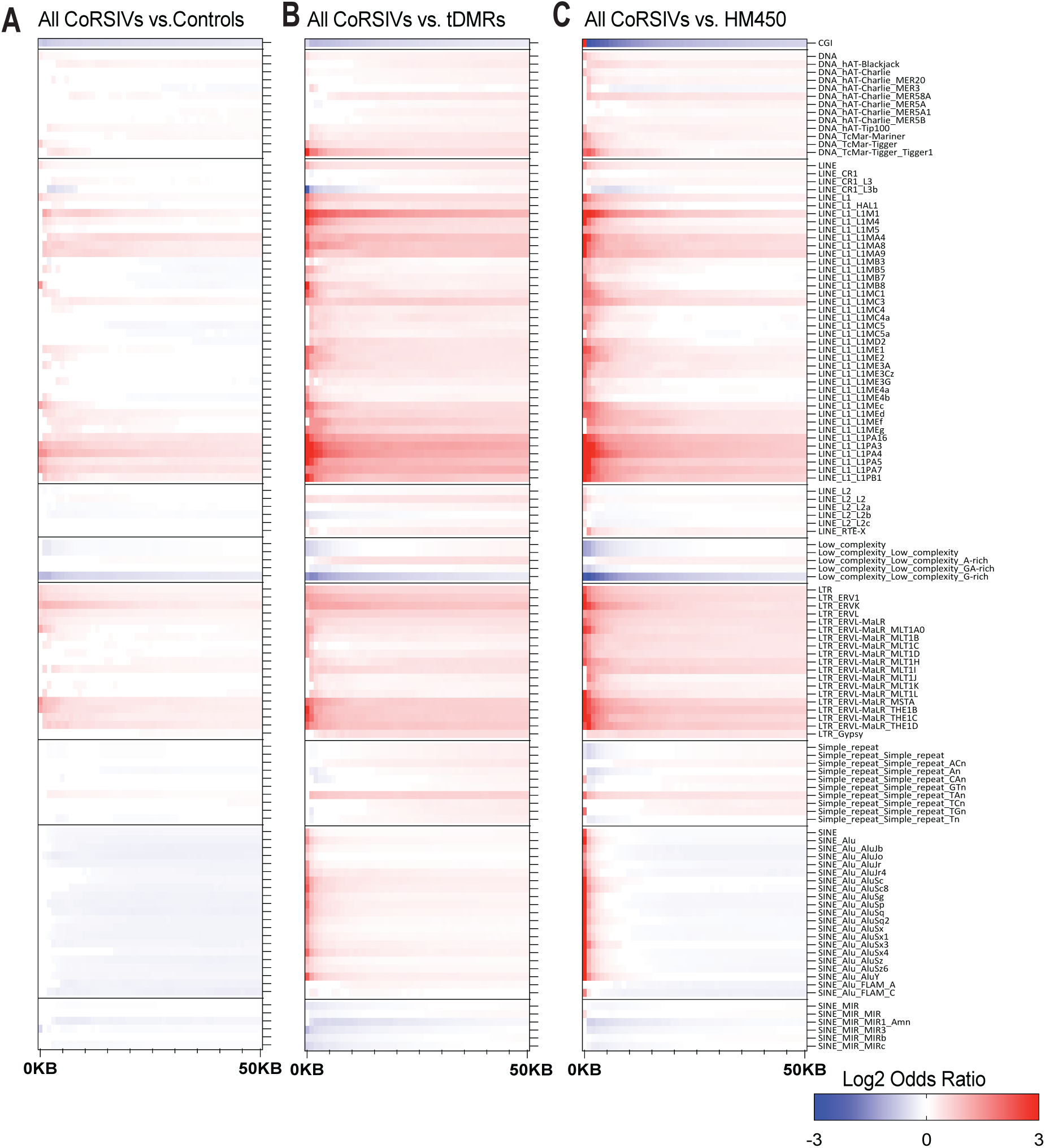
Transposable element enrichment for (A) All CoRSIVs vs. controls, (B) All CoRSIVs vs. tDMRs, and (C) All CoRSIVs vs. HM450.

**Fig. S22.**
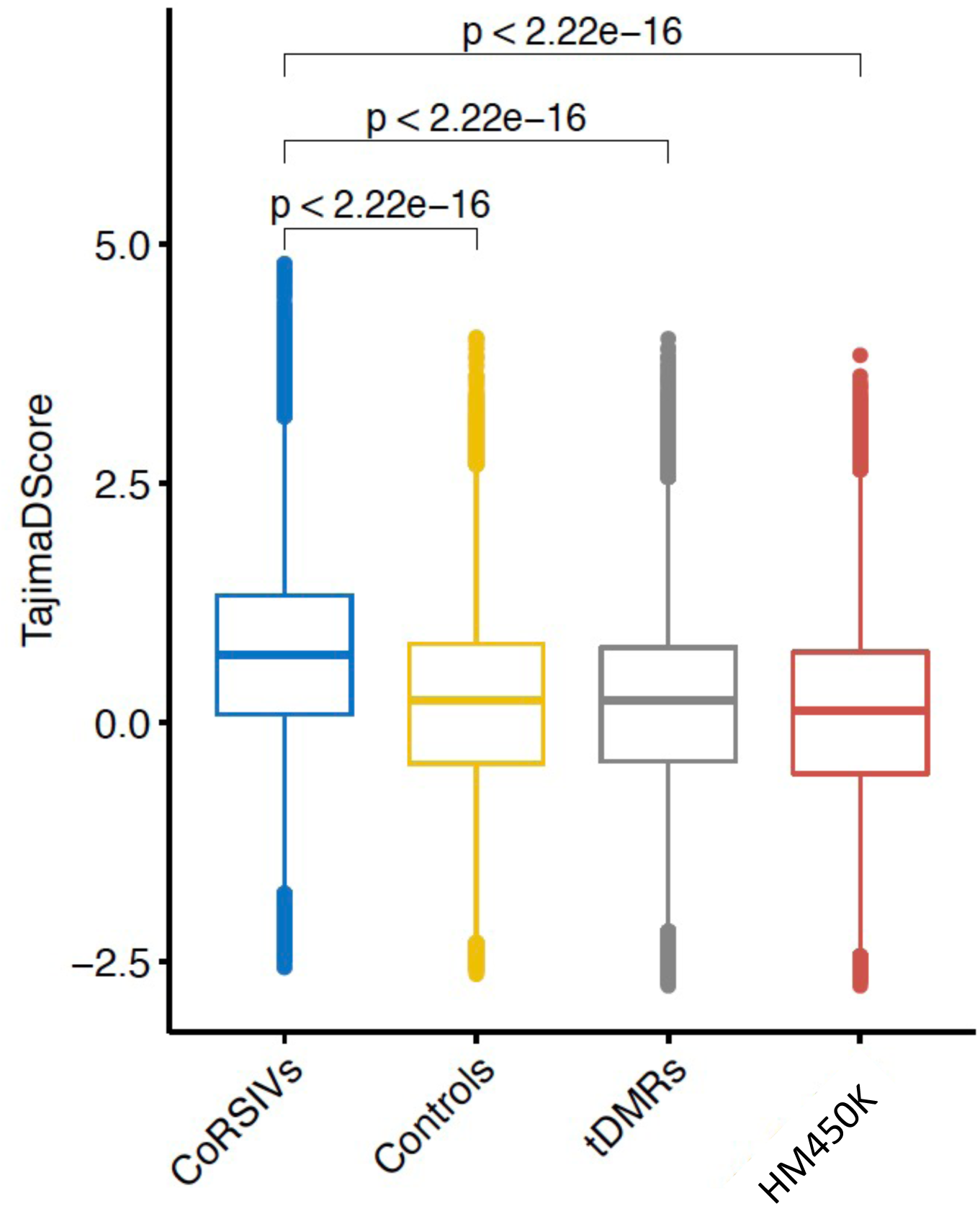
Tajima’s D Score distributions for CoRSIVs, Controls, tDMRs, and HM450 probes. Tajima’s D is higher in CoRSIVs than in the other regions, providing evidence of evolutionary selection.

**Fig. S23.**
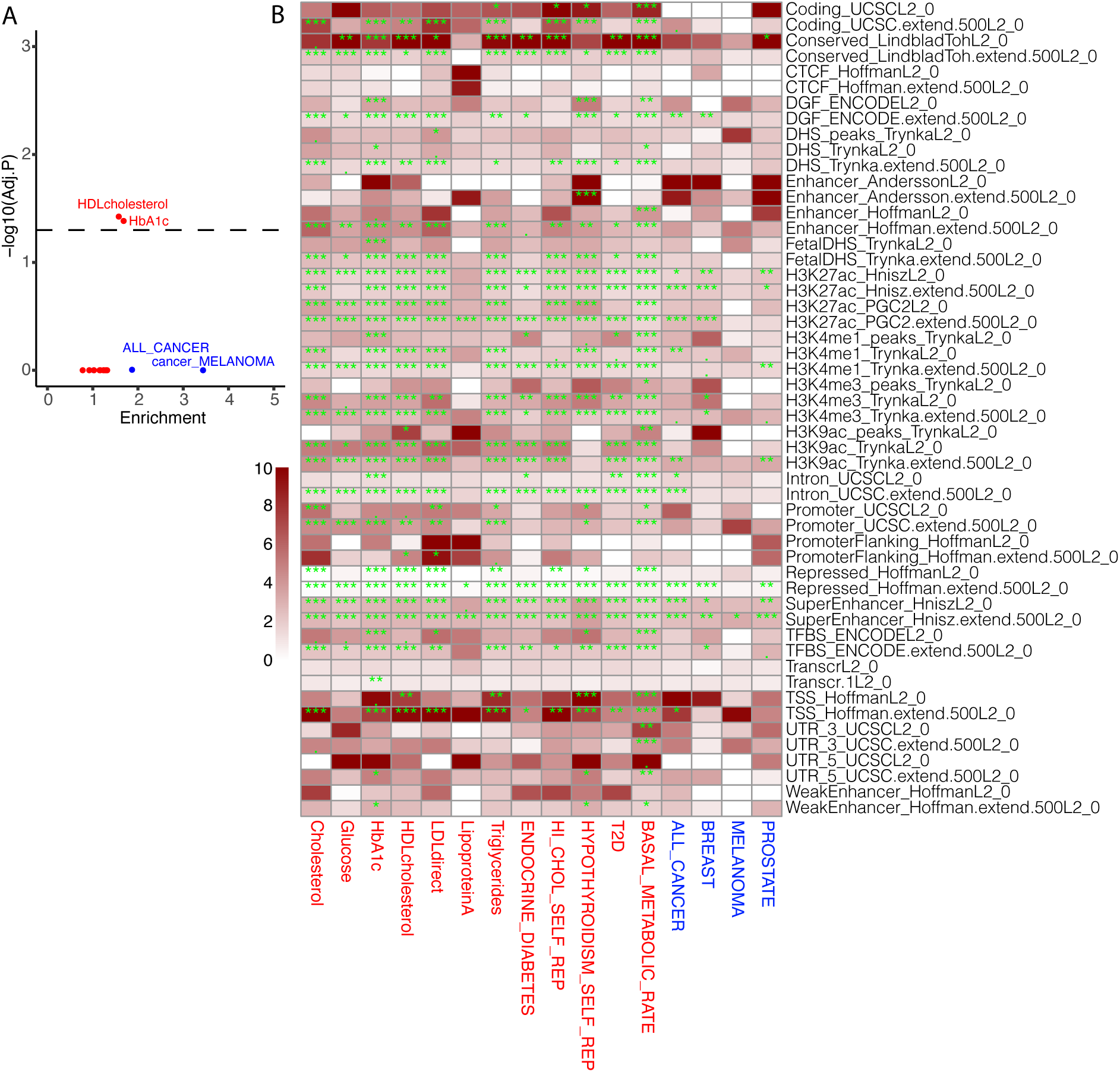
LDSC heritability enrichment score for SNVs in CoRSIV +/- 20kb, when 53 baseline features are included in the model. **(A)** LDSC enrichment score vs. Bonferroni adjusted p-value in -log10 scale for 12 metabolic traits and 4 cancer outcomes when CoRSIV +/- 20kb region and full ‘baseline’ features including 53 sequence and epigenomic features are included in the models. **(B)** LDSC Enrichment and Bonferroni adjusted p-value (green color) for 53 baseline sequence and epigenomic features when used in the models with CoRSIV +/- 20kb SNVs. 0 - 0.001 = ’***’, 0.001 - 0.01 = ’**’, 0.01 - 0.05 = ’*’, 0.05 - 0.1 = ’.’, 0.1 - 1.0 = ‘ ’.

